# Dormancy stabilizes structured food webs

**DOI:** 10.64898/2026.05.28.728563

**Authors:** Zachary R. Miller, David A. Vasseur, Pincelli M. Hull

## Abstract

Theory predicts that strong species interactions drive ecological instability, but strong interactions are common in ecosystems while strong instability appears rare. This discrepancy motivates enduring interest in ecological mechanisms that limit or counteract instability. Dormancy – reversible metabolic suppression – may be one. Dormancy is a ubiquitous life history trait found in organisms ranging from bacteria to trees. Dormant individuals form “seed banks” that are temporarily disengaged from demographic processes and species interactions, creating a memory of past ecological dynamics. Seed banks can stabilize predator-prey interactions, but whether, when, and how they affect the stability of larger ecological networks is uncertain. We show that dormancy stabilizes oscillatory dynamics in a minimal mathematical model and illustrate how dormancy converts high oscillation frequency into strong restoring force. We find that dormancy can have qualitative stabilizing effects in structured food webs that undergo Hopf bifurcations and exhibit oscillatory instability, but not in unstructured networks or those dominated by competitive or mutualistic interactions. This classification remains accurate when only a subset of species go dormant and drive stabilization. Our results clarify when dormancy can promote stability, indicating that dormancy may be an important but overlooked stabilizing factor in food webs.

## Introduction

Many ecosystems exhibit remarkable stability in the face of perturbations [1, 2, 3, 4]; yet, others show signs of instability, such as population fluctuations or alternative stable states [1, 5, 3, 6]. Understanding the balance between stabilizing and destabilizing mechanisms in large, complex ecosystems is a long-standing challenge for ecology [5, 7, 8]. Theory predicts that strong species interactions generically drive unstable dynamics [9, 10, 11, 12, 3], but factors such as population structure [13], interaction network architecture [14, 7, 8, 15], and spatial dispersal [16] can modulate or suppress instability.

Here, we explore how another fundamental but less-studied process – dormancy – shapes the stability of ecosystems. Across all major clades of life, organisms use dormancy to reduce their energetic demands and limit sensitivity or exposure to stressors [17, 18, 19, 20]. Dormancy encompasses a wide range of physiological and behavioral adaptations for reversible metabolic suppression. These adaptions exert a variety of stabilizing effects on ecological and evolutionary dynamics [19, 21, 22]. Subpopulations of dormant individuals, often called “seed banks”, can promote the maintenance of genetic diversity within populations [23], reduce temporal variation in fitness by acting as a bet-hedging strategy in variable environments [24, 18, 25], and suppress population fluctuations driven by strong density-dependence [26].

The ubiquity and steadying effects of dormancy suggest that it might also promote stability at the ecosystem level. Indeed, dormancy can stabilize the dynamics of predators and prey in models [27, 28, 29] and experimental microcosms [30] by dampening predator-prey oscillations. However, it remains unclear if and when the effects of dormancy matter in larger food webs or other ecological networks, particularly when instability is driven by collective dynamics rather than specific strong interactions. This question was studied by Hadeler and colleagues [31, 32, 33, 34, 35], who derived algebraic conditions for dormancy to stabilize an equilibrium in simple models. While these conditions apply to networks of any size or structure, they are challenging to interpret and provide little direct ecological insight, leading to very few applications (but see [27]).

Here, we develop a more general picture of when dormancy can and cannot stabilize ecological dynamics. First, we show how the algebraic conditions derived by Hadeler can be interpreted geometrically, providing new conceptual insight into the mechanism by which dormancy imparts stability. We then use the connection between an ecological network’s structure (type and organization of interactions), spectrum (eigenvalues of the community matrix), and dynamics (how instability arises) to show that the stabilizing effect of dormancy in larger ecosystems is specific: dormancy readily stabilizes structured food webs, but not most other kinds of networks. We also show that this classification remains informative in more complex models where species differ in their capacity for dormancy, suggesting unexpected simplicity in the effects of dormancy on ecological dynamics.

## Results

### Model and Stability Analysis

Following Hadeler [31, 32, 33, 34, 35], we consider a very general ecological model with a simple form of dormancy:

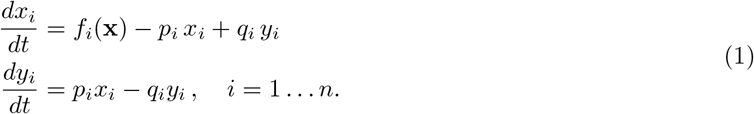

Here *x*_*i*_ is the abundance or biomass of the active individuals of species *i*, and *y*_*i*_ is the abundance of dormant individuals (the seed bank). The underlying dynamics of the community – without dormancy – are defined by the functions *f*_*i*_. These dynamics are left essentially arbitrary, but they depend only on the active populations **x** = (*x*_1_, *x*_2_, …, *x*_*n*_), implying that active individuals do not interact with dormant ones.

We assume that dormant individuals undergo no growth or mortality. Active individuals go dormant at an “inactivation” rate *p*_*i*_ ≥ 0 and dormant individuals reenter the active population at an “activation” rate *q*_*i*_ ≥ 0. This models an idealized scenario where entry and exit from dormancy are not tied to external cues, as in stochastic dormancy dynamics observed in some plants and bacteria [36, 37, 25] as a bet-hedging strategy [24, 21].

We are interested in cases where the underlying dynamics 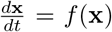 have an equilibrium **x**^⋆^ (with all 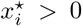) that is unstable in the absence of dormancy. Defining the community matrix 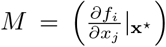, the equilibrium **x**^⋆^ is unstable if any eigenvalues of *M* have a positive real part [38]. Geometrically, these destabilizing eigenvalues of *M* lie in the right-half of the complex plane.

Adding dormancy to the model has no effect on the equilibrium abundances: **x**^⋆^ remains an equilibrium of Eq. 1 with appropriate **y**^⋆^ (Materials and Methods). However, the stability of this equilibrium may change qualitatively. To understand when and how this occurs, we start by assuming that all species enter and exit dormancy at the same rates (*p*_*i*_ = *p* and *q*_*i*_ = *q* for all *i*). We return to the more general case below.

Under this simplifying assumption, Hadeler [32] showed that if every destabilizing eigenvalue of *M* lies within a region sufficiently close to the imaginary axis, then **x**^⋆^ becomes (asymptotically) stable. In other words, dormancy extends the stability domain into the right-half of the complex plane.

We derive this augmented stability domain (ASD) in the Materials and Methods. The ASD boundary runs through the origin with vertical asymptotes at Re = *p*. The precise shape of the ASD depends on the rates *p* and *q*, as we explore below. Qualitatively, the ASD takes the form of a vertical strip between Re = 0 and Re = *p*, excluding an “indent” around the real axis. A typical ASD is shown in Fig. 1A. If all destabilizing eigenvalues of *M* (those with positive real part) fall in the shaded region, then dormancy stabilizes an otherwise unstable equilibrium.

**Figure 1.**
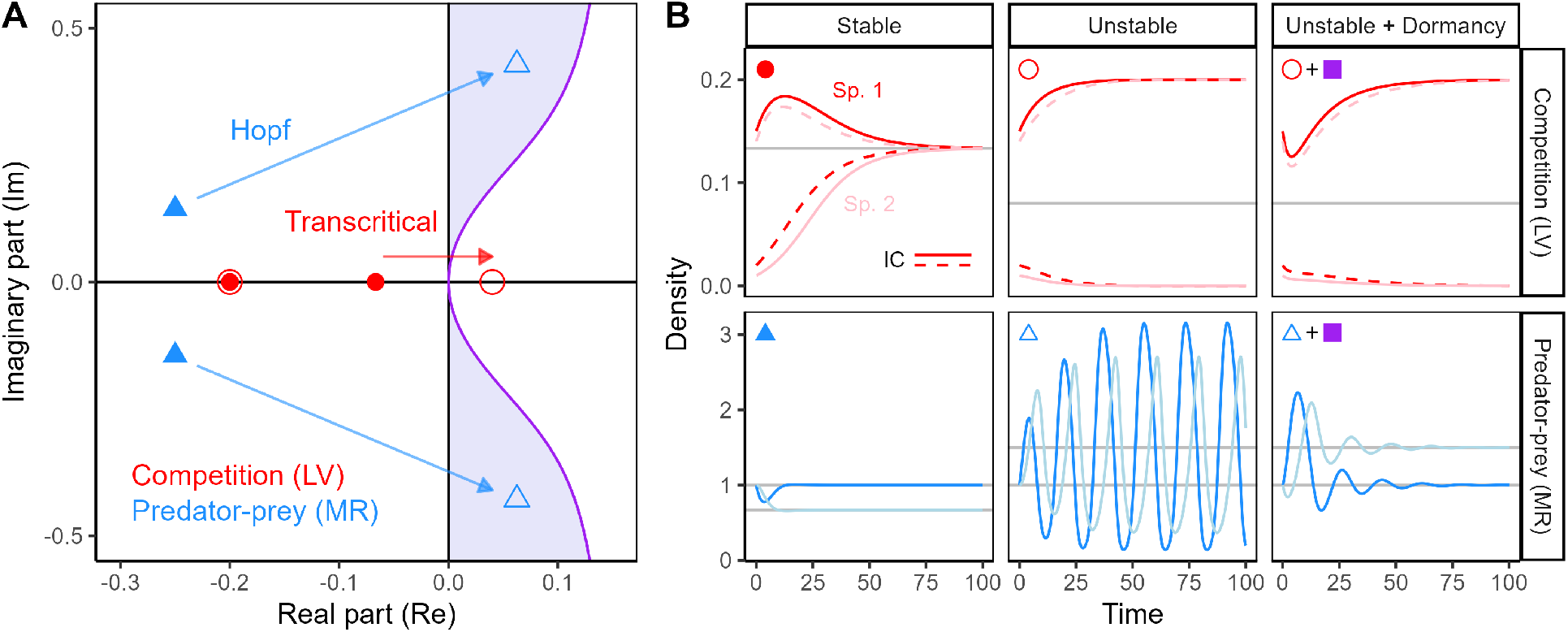
Effect of dormancy in two-species models. (A) Community matrix eigenvalues corresponding to stable (solid shapes) and unstable (open shapes) parameterizations for the Lotka-Volterra (LV) competition model (red circles) and the MacArthur-Rosenzweig (MR) predator-prey model (blue triangles). Eigenvalues in the right-half of the complex plane (Re > 0) indicate instability of the associated equilibrium. The LV model loses stability through a transcritical bifurcation as a real eigenvalue passes through the origin; the MR model loses stability through a Hopf bifurcation as a pair of complex eigenvalues crosses the imaginary axis. The augmented stability domain (ASD) due to dormancy is shown in purple. (B) Simulated dynamics for stable and unstable parameterizations. Rightmost column shows the unstable parameterization when dormancy is added to the model. Shading indicates species identity; line type indicates different initial conditions (LV only). The unstable LV model exhibits a priority effect that is not stabilized by dormancy. The unstable MR model exhibits oscillations, which are stabilized. LV model parameters: *r* = 0.2, *α* = 0.5 (stable) and 1.5 (unstable). MR model parameters: *µ* = 0.5, *K* = 1.5 (stable) and 4 (unstable). Dormancy parameters: *p* = 0.2, *q* = 0.15 for both models.

The shape of the ASD means that dormancy cannot stabilize systems whose eigenvalues have small imaginary parts or real parts greater than *p* [35]. In particular, purely real eigenvalues can never be stabilized. On the other hand, if all destabilizing eigenvalues have large imaginary parts, dormancy is likely to be stabilizing. In the next two sections, we relate this characterization of the ASD to the nature of the ecological dynamics.

### Two-species models

We first compare the effect of dormancy in two classical two-species models. We consider a model of competition, the symmetric Lotka-Volterra (LV) model [39], with carrying capacity *K* = 1 and interspecific competition strength *α*. For *α <* 1 (relatively weak interspecific competition), the LV model has a stable coexistence equilibrium, indicated by two real, negative eigenvalues of *M* (solid red points in Fig. 1A). As *α* increases, one eigenvalue slides along the real axis and eventually becomes positive for *α >* 1, where competition is stronger between than within species (open red points). The coexistence equilibrium is now unstable (Fig. 1B, top), and the model exhibits a priority effect (alternative stable states). This situation, where stability is lost as a real eigenvalue passes through the origin and becomes positive, is called a transcritical bifurcation [38].

Dormancy cannot stabilize this type of bifurcation. Because the destabilizing eigenvalue is real, it falls outside of the ASD (Fig. 1A). As a result, adding dormancy to the model has little effect – we still find a priority effect due to the unstable equilibrium (Fig. 1B, right).

We also consider a predator-prey model, the MacArthur-Rosenzweig (MR) model [40, 27]. In the MR model, a predator with density-independent mortality consumes a logistically-growing prey via a saturating functional response. If the prey’s carrying capacity *K* exceeds a critical threshold, the predator-prey equilibrium becomes unstable, a phenomenon known as the “paradox of enrichment” [40]. The eigenvalues of the community matrix are complex, and as *K* increases, their real parts shift from negative (stable) to positive (unstable) while the imaginary parts remain nonzero (solid and open blue triangles in Fig. 1A, respectively). This is a Hopf bifurcation [38], associated with oscillatory behavior around the equilibrium (Fig. 1B, bottom). Note that for ecological models, where the elements of *M* are real numbers, any complex eigenvalues come in conjugate pairs (i.e., *λ* = *α* ± *βi*), so a Hopf bifurcation occurs as a pair of eigenvalues crosses the imaginary axis.

In this case, adding dormancy to the model has a more dramatic effect. In Fig. 1A, we see that the eigenvalues of the unstable MR model fall within the ASD. Accordingly, with dormancy, the oscillations are strongly damped, and the dynamics quickly settle into a stable equilibrium (Fig. 1B, right) [27].

Both of these models exhibit instability, indicated by eigenvalues with positive real parts, but they lose stability in fundamentally different ways. We see that the effect of dormancy is mediated by the type of bifurcation: dormancy is uniquely stabilizing against Hopf bifurcations and associated oscillatory instability [31, 32, 33, 35].

### Phase-plane geometry and the origins of stabilization

Why can dormancy stabilize oscillatory dynamics but not other kinds of unstable dynamics? The ASD tells us when but not why to expect stabilization, so we turn to a closer examination of the MR model dynamics for more insight. Fig. 2A shows these dynamics in the prey-predator plane. We consider a scenario where the system is initially sitting at a stable equilibrium (black point) before the prey’s carrying capacity is suddenly increased. Without dormancy (blue trajectory), the dynamics spiral outward and approach a limit cycle. For the same model with dormancy (purple trajectory), the dynamics instead spiral inward to a new stable equilibrium point.

**Figure 2.**
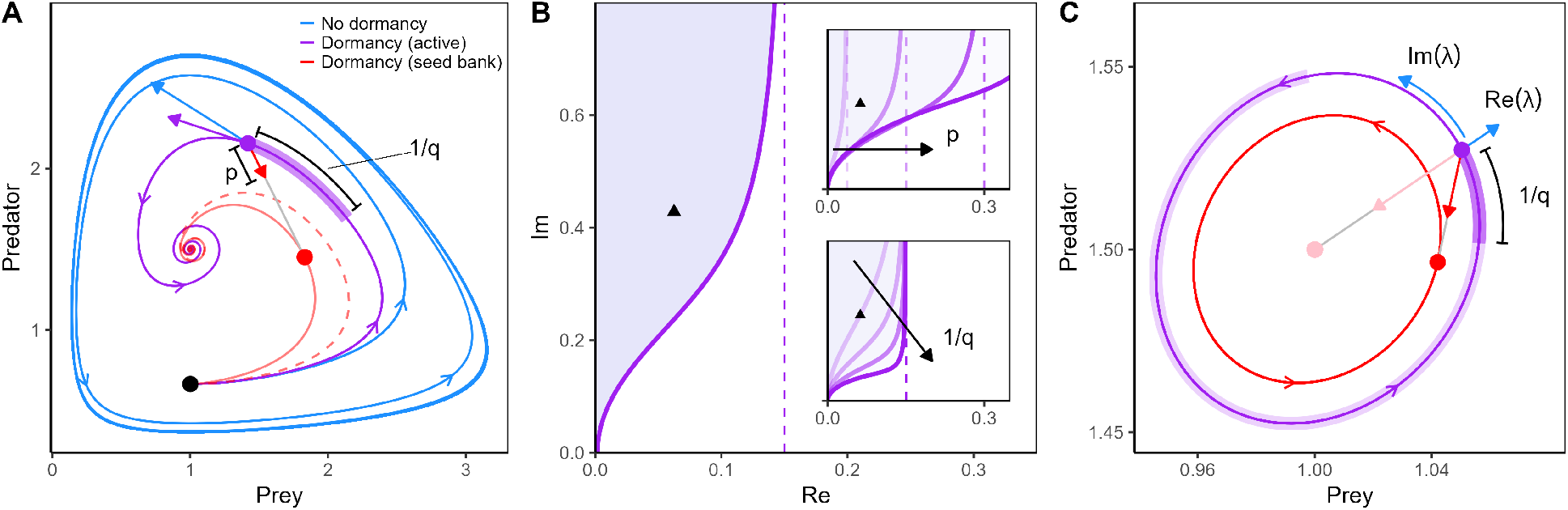
Stabilizing effect of dormancy in a predator-prey model. (A) MR model dynamics with (purple/red) and without (blue) dormancy, plotted in the predator-prey plane. Both systems start at a stable equilibrium with *K* = 1.5 (black point); when the prey’s carrying capacity is increased to *K* = 4, the system without dormancy spirals outward to a limit cycle, while the system with dormancy converges to a new stable equilibrium. Active dynamics (purple trajectory) are influenced by the underlying dynamics (blue arrow) and the pull of dormancy (red arrow) toward the seed bank state (red point), which converges to a weighted time-average of the active state (red dashed line). Parameters *p* and *q* control the magnitude and “memory” of the seed bank effect, respectively. (B) shows how these parameters affect the ASD (shaded region): increasing *p* shifts the vertical asymptote (top inset), while decreasing *q* (increasing 1*/q*) pulls the ASD boundary downward (bottom inset), stabilizing eigenvalues with smaller imaginary parts. Because the ASD and eigenvalues (black triangles) are symmetric over the real axis, we only plot Im > 0. (C) When dormancy is marginally stabilizing (i.e., eigenvalues fall on the ASD boundary), the dynamics oscillate neutrally. Close to equilibrium, the underlying dynamics (blue) can be decomposed into a rotational part proportional to |Im(*λ*) | and a radial (outward) component proportional to Re(*λ*). The seed bank (red) also oscillates, lagging behind the active state. Decreasing *q* lengthens the seed bank memory and shrinks the seed bank orbit, producing a more inward (stabilizing) pull on the dynamics (pink).

With dormancy, the active dynamics can be decomposed into a component due to the underlying model *f* (**x**) (blue) and a component due to dormancy (red), according to Eq. 1. In Fig. 2A, we visualize this decomposition at one point in time (arrows). The effect of dormancy depends on the composition of the seed bank, which converges to a time-average of the active state, weighted toward more recent times (red dashed line in Fig. 2A). Intuitively, the abundance of each species in the seed bank reflects its abundance in the active community at earlier times. The model dynamics can therefore be approximated (Materials and Methods) as

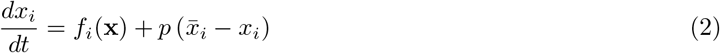

where 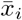 is the weighted time-average 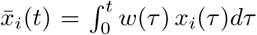 with the time-weighting function *w*(*τ*) proportional to exp(*qτ*).

From Eq. 2, we see that dormancy “pulls” the active dynamics toward the averaged historical state (red arrow in Fig. 2A). For dynamics that oscillate in a convex orbit around an unstable equilibrium, as in a predator-prey cycle, the time-average of the dynamics, 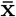, must lie inside the orbit. Thus, dormancy pulls the dynamics inward. If this effect is (on average) sufficiently strong compared to the outward pull of the underlying dynamics (blue arrow), then trajectories will spiral inward to the equilibrium. For non-oscillatory dynamics, such as the priority effect in the LV model, this is not the case; trajectories move away from the equilibrium, but not around it, and consequently the time-average of the active state does not converge toward the equilibrium.

This geometric view provides additional insight into the effect of the rates *p* and *q* on the ASD (Fig. 2B). In Eq. 2, the inactivation rate, *p*, scales the relative contribution of dormancy to the net dynamics. Larger *p* allows dormancy to overcome stronger underlying instability, indicated by the real part of the destabilizing eigenvalue(s). Increasing *p* therefore shifts the vertical asymptote of the ASD to the right (Fig. 2B, top inset).

The activation rate, *q*, controls the weighting function, *w*, in the time-average. Larger *q* places more weight on the recent past, resulting in a seed bank with a shorter “memory”. Intuitively, this is because 1*/q* sets the mean time spent in dormancy. Smaller *q* produces a flatter weighting of past states, leading the seed bank state to converge more strongly toward the equilibrium. Fig. 2B (bottom inset) shows that decreasing *q* (increasing 1*/q*) pulls the boundary of the ASD toward the real axis. This allows dormancy to stabilize dynamics whose eigenvalues have smaller imaginary parts, which control the frequency of oscillations. For dormancy to generate a sufficiently stabilizing inward pull, the seed bank must produce sufficiently strong time-averaging, which is enhanced by both a long seed bank memory (low *q*) and higher-frequency dynamics (associated with large Im).

This account becomes precise when the system’s eigenvalues, *λ*_±_, fall exactly on the boundary of the ASD. In Fig. 2C, we plot the dynamics for the same MR model as in panel A, but with *q* adjusted to produce a marginally stable equilibrium. The system now undergoes small oscillations around equilibrium, with the seed bank lagging behind the active community state. Close to equilibrium, the underlying dynamics (blue arrows) decompose into a rotational component, proportional to |Im(*λ*)|, and a radial (outward) component, proportional to Re(*λ*). The inward pull generated by dormancy is determined by the projection of the dormancy effect (red arrow) onto the radial direction. For this choice of parameters, the inward pull of dormancy and the outward pull of the underlying dynamics are precisely balanced. Any increase in *p* would increase the magnitude of the dormancy effect, pulling the dynamics inward. Similarly, any decrease in *q* would lengthen the memory of the seed bank, causing its orbit (red) to shrink and exert a more inward pull on the dynamics. An equivalent effect would come from increasing the rotational speed, Im(*λ*), effectively lengthening the seed bank memory by increasing the frequency of oscillations. For very small *q* or large |Im(*λ*) |, the seed bank orbit shrinks to a point, and the pull of dormancy, with magnitude *p*, is directly opposed to the outward pull with magnitude Re(*λ*). In this configuration (pink), the seed bank effect on the dynamics is maximized, explaining the vertical asymptote in the ASD.

### Multispecies models

For two-species models, there is a tight connection between the type of interaction and the type of instability. Competitive (−, −) or mutualistic (+, +) systems have real eigenvalues and lose stability via transcritical bifurcations (SI Section 3A). Only predator-prey (+,−) pairs can undergo Hopf bifurcations, leading to oscillatory dynamics that can be stabilized by dormancy.

With more than two species, however, the relationship between ecological interactions and eigenvalues is much more complex. It is affected by both the type of interactions in the community and their organization. To investigate when dormancy can stabilize the dynamics of larger communities, we study random and idealized community matrices, for which we can link properties of the ecological network to the distribution of its eigenvalues. Stability is determined by the interplay of the eigenvalue and ASD geometries (Fig. 3).

**Figure 3.**
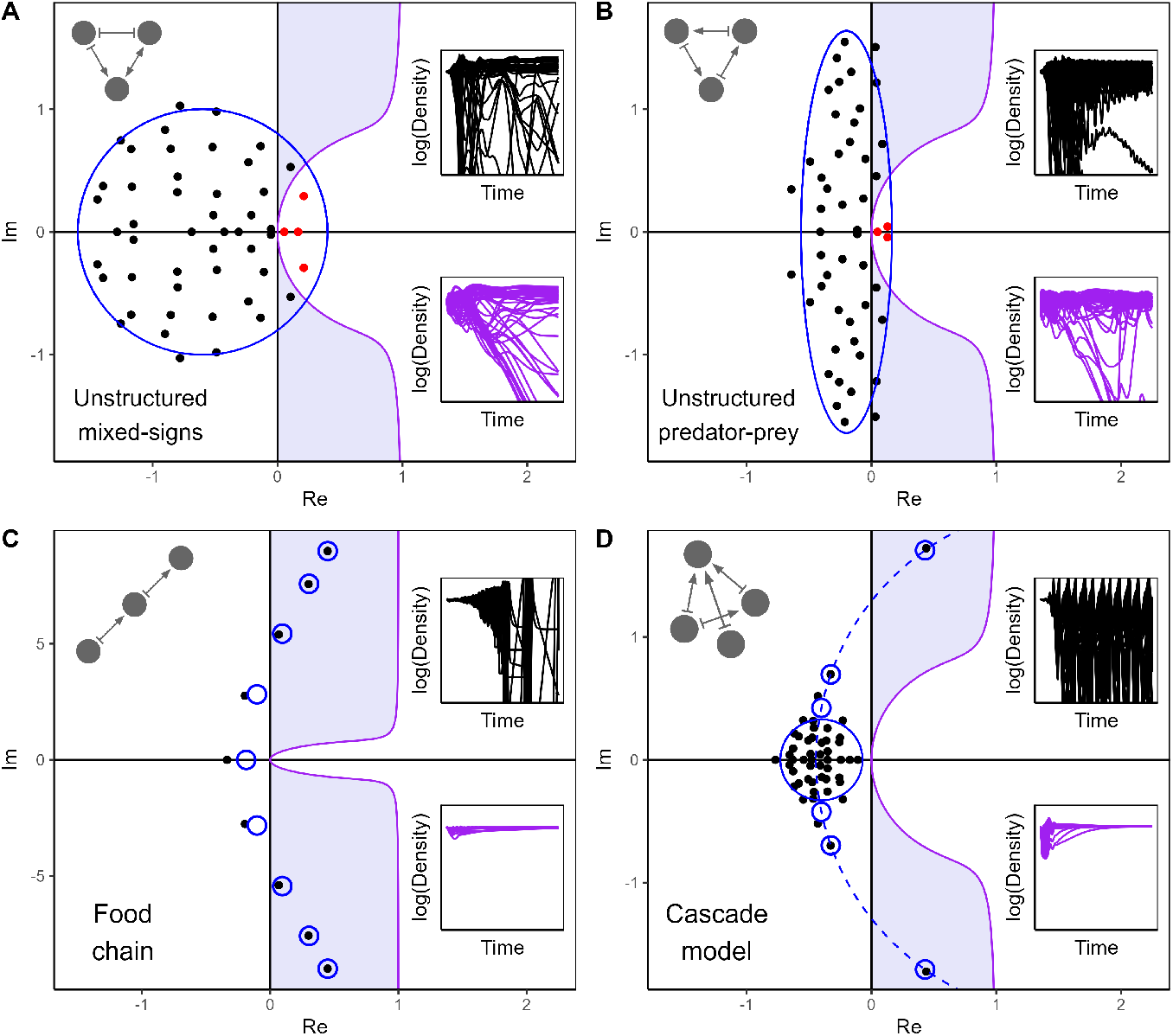
Effect of dormancy in multispecies models. Community matrix eigenvalues are shown for different network models in relation to an example ASD (purple shaded region), indicating the stabilizing effect of dormancy. Theoretical eigenvalue distributions are shown in blue; points show eigenvalues for one model realization. Eigenvalues that fall to the right of the ASD (red) remain destabilizing with dormancy. For each model, simulated dynamics without (black) and with (purple) dormancy are shown in inset panels. Dormancy is not stabilizing for unstructured mixed (A) or predator-prey interactions (B). Stability in the food chain (C) and structured (cascade model) food web (D) is controlled by eigenvalues with large imaginary parts; consequently, these dynamics are stabilized by dormancy. Dynamics were generated by numerical integration of Eq. 1, using generalized Lotka-Volterra models with the desired community matrix at a feasible (strictly positive) equilibrium (see SI Section 5 for details). *n* = 50 species in all panels except C, where *n* = 9. Dormancy parameters: *p* = 1 and *q* = 0.25 for all models. Note the distinct y-axis scale in panel C.

First, we consider an unstructured network where the effect of each species on every other at equilibrium, *m*_*ij*_, is an independent, identically-distributed (*iid*) random variable with mean zero. May [9] famously used this type of random matrix as a simple model for large, complex ecosystems. The community matrix eigenvalues in this case are uniformly distributed over a circle in the complex plane, centered on the real line – a result known as the Circular Law [7, 41]. The circle’s center point is determined by the strength of self-regulation (diagonal coefficients in the community matrix). If self-regulation is sufficiently weak relative to the variance in interaction strengths (controlling the radius), the circle extends into the right-half plane, producing instability. As Fig. 3A illustrates, this eigenvalue geometry ensures that the rightmost eigenvalues – those with the largest real parts, which control stability – will usually have small imaginary parts. In large networks, this becomes essentially guaranteed. These eigenvalues fall outside of the ASD, so dormancy cannot stabilize the equilibrium.

In May’s random matrix ensemble, *m*_*ij*_ and *m*_*ji*_ are sampled independently, so all combinations of interaction signs, corresponding to competitive, mutualistic, and predator-prey interactions, are possible. Because dormancy stabilizes predator-prey pairs in isolation, we might expect a stabilizing effect in networks composed exclusively of predator-prey (+, −) interactions. As a simple variation on the mixed-interaction network, we can consider a community matrix where all pairs *m*_*ij*_ and *m*_*ji*_ have opposite signs. We randomly assign the direction of the interaction (i.e., whether *i* or *j* is the prey) independently for each pair, and sample all interaction magnitudes |*m*_*ij*_| *iid*. This type of community matrix models trophic interactions, but not realistic trophic structure. For example, this procedure generates many trophic cycles, i.e., closed chains of predators and prey.

Allesina and Tang [7] showed that the eigenvalues for this random predator-prey network are uniformly distributed over an ellipse, generalizing the Circular Law (Fig. 3B). The negative correlation between *m*_*ij*_ and *m*_*ji*_ produces an elliptical distribution that is wider in the imaginary direction and compressed along the real axis. While this leads to more eigenvalue pairs with large imaginary parts, the elliptical shape ensures that at least some of the rightmost eigenvalues have small imaginary parts with high probability. Consequently, it is again unlikely that dormancy can stabilize this type of network. Fig. 3B shows that, as for the mixed-interaction network, the presence of destabilizing eigenvalues near the real line leads to complex, fluctuating dynamics and species extinctions, with (purple) or without (black) dormancy.

This example reveals that the stabilizing effect of dormancy on predator-prey interactions does not automatically extend to larger trophic networks. However, trophic interactions in nature are not arranged randomly. Most notably, empirical food webs exhibit trophic hierarchy, with few or no trophic cycles [42, 43].

As an idealized model of *structured* predator-prey interactions, we first consider a linear food chain. The dynamics and stability of food chain models have been studied extensively [44, 11, 10], but the geometry of their eigenvalues in the complex plane has received less attention. We use a perturbation approach to approximate the community matrix eigenvalues for a simple food chain where consumption effects are symmetric at each trophic level (SI Section 1). Under fairly general conditions where there is enhanced self-regulation in the basal species (e.g., carrying capacity) and/or apex predator, the eigenvalues of the community matrix lie on an arc opening to the right in the complex plane (Fig. 3C). The rightmost eigenvalues have large imaginary parts, while those close to the real line have more negative real parts. This eigenvalue geometry readily produces unstable dynamics that can be stabilized by dormancy.

Finally, to incorporate trophic structure in a more realistic way, we consider community matrices structured by the cascade model for food webs [42, 45], where species are ordered from basal to apex and each species can only consume those lower in the trophic ranking. Corresponding to this food web model, Allesina et al. [46] studied random matrices where the elements *m*_*ij*_ are sampled independently but with distinct distributions above and below the diagonal, such that *m*_*ij*_ < 0 for *i* < *j* (effects of predators on prey) and *m*_*ij*_ > 0 for *i* > *j* (effects of prey on predators). In this case, the set of eigenvalues has a more complex structure: the majority fall into an elliptical “bulk”, as in the unstructured cases, but a few “outlier” eigenvalues are distributed along an arc that passes through the bulk (Fig. 3D). The orientation of the outlier arc depends on the ratio of the mean prey → predator and predator → prey effects at equilibrium. When the mean effect of prey on predators (positive *m*_*ij*_ elements) is larger than the mean effect of predators on prey (negative *m*_*ij*_), the outlier arc opens rightward, as for the linear food chain. This corresponds to an ecological scenario where donor-control predominates in the food web [47, 14, 45].

In this case, the rightmost eigenvalues in the outlier arc have large imaginary parts. In order for these eigenvalues to control stability, the elliptical bulk must also be sufficiently small relative to the outlier arc. Qualitatively, the size of the bulk is controlled by the variance in interaction strengths, while the size of the arc is controlled by the means (Materials and Methods). If the variance is sufficiently small relative to the means, indicating that the structure in the matrix dominates over the noise in individual interactions, then the rightmost eigenvalues are outliers. As Fig. 3D shows, this produces an eigenvalue geometry that is stabilized by dormancy.

These two examples – a simple food chain and a random food web – show that dormancy can stabilize large trophic networks, but only when accounting for network structure. When interactions are organized in a trophic hierarchy, as in empirical food webs, the nature of instability changes – stability is determined by eigenvalues with large imaginary parts. Correspondingly, the unstable dynamics in these cases (without dormancy) are strongly oscillatory, particularly in Fig. 3D. However, with many species, it is not straightforward to predict the effect of dormancy directly from the dynamics. The unstructured predator-prey dynamics (Fig. 3B) also exhibit oscillatory behavior, but these oscillatory modes occur along with non-oscillatory instability that cannot be stabilized by dormancy. To distinguish these cases, it is necessary to resolve the eigenvalue geometry of the network.

Crucially, it is the combination of real and imaginary eigenvalue components that determine the effect of dormancy. Focusing on real parts alone – a common measure of (in)stability in ecological models [48, 46] – would suggest that the two structured models in Fig. 3 (C and D) are more unstable, and thus more difficult to stabilize, than the unstructured models (e.g., via self-regulation). Yet, when stabilization originates from dormancy, the opposite is true.

### Rate heterogeneity

Our results so far rely on the simplifying assumption that all species enter and exit dormancy at the same rates (i.e., homogeneous rates), which is necessary to derive the ASD. This assumption is unrealistic, especially in large food webs where species will have distinct life histories and only some species are likely to go dormant at all. When (in)activation rates vary between species (including zero values for those without dormancy), it is much more challenging to mathematically characterize the effect of dormancy on stability. Dormancy can even sometimes destabilize an otherwise stable equilibrium, which is not possible when all rates are the same [35].

Yet, we hypothesize that our main conclusions from the homogeneous case carry over to the heterogeneous one, at least qualitatively. We provide two lines of evidence: First, we prove in SI Section 3A that if the underlying community matrix *M* is symmetric (more generally, *symmetrizable*), then dormancy can never be stabilizing, even with heterogeneous rates. Symmetry requires that all interactions are either competitive or mutualistic, consistent with our expectation that dormancy generally does not stabilize non-trophic interactions. All 2×2 matrices with *m*_12_*m*_21_ > 0 are symmetrizable, so this result implies that dormancy can never stabilize a competitive (−, −) or mutualistic (+, +) pair.

Second, we use a numerical approach to explore the effect of dormancy when only some species go dormant. Using the food web model described above, we construct community matrices in three parameter regimes [45, 46]: donor-controlled interactions (D) where the mean effect of prey on predators is twice the mean effect of predators on prey, recipient-controlled interactions (R) where the mean effects are reversed, and balanced interactions (B) where these effects are equal on average. Eigenvalue distributions for these three parameterizations are plotted in Fig. 4 (top left). In each regime there is a bulk of eigenvalues and a pair of outliers distinct from the bulk. As expected, these outliers fall to the right of the bulk for the donor-controlled regime, but not for the other two. Thus, with homogeneous rates, we would predict that only the donor-controlled parametrization can be stabilized.

**Figure 4.**
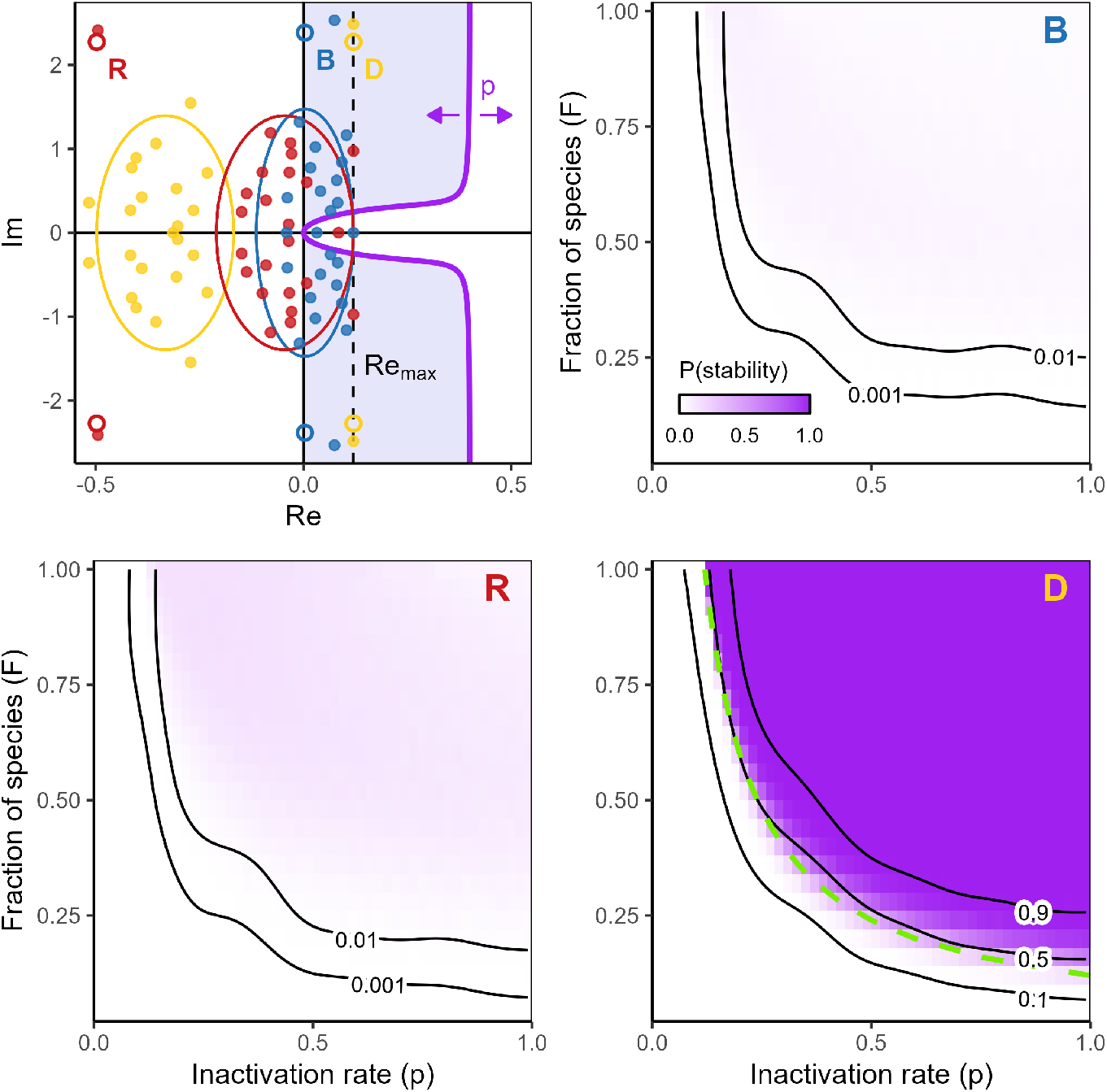
Dormancy stabilizes structured food webs when only some species go dormant. We plot the probability of stability in three parameter regimes (B, R, D) of the cascade food web model, as a function of the fraction of species with dormancy, *F* (y-axis), and inactivation rate, *p* (x-axis). White indicates zero probability of stability across 100 random realizations of the food web and 25 random species subsets for each web; dark purple indicates stabilization in all 2500 web-subset combinations. Top left panel shows the theoretical eigenvalue distributions for the three parameterizations in relation to the ASD (purple shaded region), visualized for *p* = 0.4. Points are eigenvalues for one community matrix realization. In the donor-controlled parameter regime (D, yellow), stability is controlled by outlier eigenvalues with large imaginary parts, leading to a high probability of stability whenever the product of *p* and *F* exceeds the real part of these eigenvalues (*pF* = Re_max_, green dashed curve). In the recipient-controlled (R, red) and balanced (B, blue) parameter regimes, stability is typically controlled by eigenvalues with small imaginary parts, making stabilization unlikely for all values of *F* and *p*. Smoothed contour lines are generated from a GAM (beta regression) fit to the probability response surface. All underlying community matrices have *n* = 25 species and Re_max_ = 0.12.

We generate 100 random food web matrices in each parameter regime, adjusting self-regulation effects (diagonal coefficients) to establish a consistent baseline instability, Re_max_(*λ*) *>* 0. To allow heterogeneity in dormancy, we set *p*_*i*_ = *p* and *q*_*i*_ = *q* in a subset of species with dormancy, and *p*_*i*_, *q*_*i*_ = 0 in all others. We vary the fraction of species that go dormant, *F*, and the inactivation rate, *p* (fixing *q* for simplicity). For every *F* - *p* combination, we select 25 distinct, random subsets of species to have dormancy and evaluate stability numerically in all 100 webs (yielding 2500 total stability assessments for each *F* - *p* pair).

Fig. 4 shows the probability of stability across *F* and *p* in the three parameter regimes (B, R, and D). In line with our expectation based on homogeneous rates, dormancy has a strong stabilizing effect across a broad range of *F* and *p* combinations in the donor-controlled regime, but with recipient-controlled or balanced interactions, the probability of stability is always low. In the donor-controlled regime, we observe a threshold-like behavior: once the product of *F* and *p* is greater than *≈*Re_max_(*λ*), which is the value of *p* where stability is gained when all species have dormancy (*F* = 1), the probability of stability jumps from near zero to nearly one.

Our numerical results suggest that the ASD is highly predictive of stability even when rate homogeneity is violated. In SI Section 3B, we show that this remains true when rates *p*_*i*_ and *q*_*i*_ vary continuously. In the donor-controlled regime, dormancy is highly stabilizing when a sufficient number of species go dormant with a sufficiently high inactivation rate. Even a small fraction of species (as low as 20% in Fig. 4) can reliably drive a qualitative change in stability if *p* is sufficiently large. In the other two parameter regimes, where stability is not controlled by outlier eigenvalues, dormancy rarely has an effect on stability. In a small fraction of food webs, we do find stabilization – these are cases where chance fluctuations in the eigenvalue distribution produce destabilizing eigenvalues with large imaginary parts (Fig. S6). Thus, these exceptions confirm the rule that dormancy is stabilizing when eigenvalues with large imaginary parts drive instability.

Interestingly, while the effect of dormancy is sensitive to the interaction network structure, it is relatively insensitive to which species in the network go dormant. In the donor-controlled regime, almost all variation in stability is explained by *F* and *p*; the identities of the species with dormancy can only matter in a narrow band around *pF ≈*Re_max_(*λ*). In this band, we find that dormancy is more stabilizing when it occurs in species that are more connected in the food web and have stronger interactions (Fig. S3-5). Perhaps surprisingly, we find no effect of trophic position (Fig. S3-5).

## Discussion

The stabilizing effect of dormancy on predator-prey interactions has been demonstrated empirically [30] and studied in some theoretical depth [27, 28, 49, 29]. Yet, if and how this effect extends to larger ecological networks has remained largely unexplored. In ecology, generalizing predictions from species pairs to the large, complex communities found in nature is an enduring challenge that often yields surprises [15, 50, 51, 52]. Here, we combined stability criteria derived by Hadeler [31, 32, 33, 34, 35] with insights on the relationship between network structure and eigenvalue geometry [7, 46] to understand how and when dormancy can stabilize multispecies ecosystems. We showed that despite the consistent stabilizing effect of dormancy in predator-prey interactions, large trophic networks are difficult to stabilize with dormancy unless they possess additional structure. The key structural feature that we identify, trophic hierarchy, is a hallmark of empirical food webs [42, 43], suggesting that dormancy might play an important stabilizing role in nature.

We also found that dormancy is unlikely to suppress instability driven by competitive or mutualistic interactions. We show in SI Section 4 that there are exceptions to this rule, such as intransitive (rock-paper-scissors) competitive loops, but it is unclear if these cases can scale up to larger competitive networks (or do in nature [53]). Additionally, these exceptions again confirm the more general rule that dormancy stabilizes oscillatory instability associated with Hopf bifurcations. By analyzing the geometry of stabilization in the predator-prey plane and linking this picture to the algebraic ASD, we provide a clearer understanding of why dormancy stabilizes against Hopf bifurcations, and how seed banks act as a mechanism to convert high oscillation frequency into reduced oscillation amplitude. This mechanistic picture helps clarify why dormancy tends to stabilize oscillatory dynamics, while time-lags, which do not generate the same consistent “pull” toward equilibrium, are often destabilizing [54, 55].

The connections we identify between network structure, eigenvalue geometry, ecological dynamics and the effect of dormancy appear to hold broadly when only some species have the capacity for dormancy, or more generally when species enter and exit dormancy at different rates (Fig. S8). This suggests that dormancy can have a strong stabilizing effect driven by relatively few species, and that the ASD, derived under highly simplifying assumptions, retains surprising predictive power in such systems. Importantly, our numerical results show that the effect of dormancy is largely independent of trophic position (Fig. S3-5). Thus, although dormancy is distributed non-randomly in natural food webs (e.g., more prevalent among basal species [19, 21, 20]), its effect on stability is unlikely to depend on the specific relationship between dormancy and trophic rank.

We demonstrated the importance of trophic hierarchy in two specific network models (food chain and cascade model), chosen for their tractability. However, Allesina et al. [46] showed that the random cascade model captures the eigenvalue distribution of the more realistic niche model [43] and empirical food webs parametrized by allometric scaling relationships. Additionally, stability is controlled by eigenvalues with large imaginary parts in other models with hierarchical structure, such as bipartite trophic networks [56]. Thus, we hypothesize that trophic hierarchy is the operative feature driving this eigenvalue geometry and associated oscillatory dynamics.

However, trophic hierarchy is not necessarily sufficient for dormancy to be stabilizing. In the random cascade model, we found that dormancy is stabilizing only when donor-control predominates; that is, when the effects of prey on predators are stronger on average than the effects of predators on prey. The prevalence of donor-controlled interactions in food webs has been enduringly and sometimes hotly debated [14, 47, 57], but donor-control can arise due to predator interference [57], nonlethal consumptive effects (e.g., many herbivores and parasites), or when predators primarily consume sick, senescent, or otherwise non-reproductive prey. Allesina et al. [46] also hypothesized that, because the elements of the community matrix are scaled by population sizes, these parameterizations may be more likely in food webs with inverted biomass pyramids, such as aquatic systems [58]. Overall, we suggest that dormancy is most likely to have an important stabilizing effect in ecosystems where (i) donor-control is likely, (ii) dormancy is prevalent, and (iii) oscillatory dynamics are documented, such as marine food webs and “non-traditional” food webs like host-pathogen systems and parasite-dominated webs [59].

Our analysis has important limitations. We assume that species enter and exit dormancy at rates that are independent of the community or environmental state. This assumption may be reasonable in some systems where environmental predictability is low and stochastic dormancy strategies are optimal [21]. In addition, as noted by Hadeler [35], if (in)activation rates are density-dependent, then equilibrium population sizes remain unaffected by dormancy, and the structure of the community matrix is unchanged, but with the values of *p*_*i*_ and *q*_*i*_ dependent on the population sizes. Thus, relaxing the homogeneous rate assumption also opens the door to studying density-dependent rates. However, this type of model still neglects age and stage structure [13], which may be important when dormancy occurs at a specific point in the life cycle. We also assume that mortality in dormancy is negligible. The consequences of this assumption are harder to constrain, because mortality in the seed bank will alter equilibrium abundances, changing the underlying community matrix. Non-negligible mortality would shorten the seed bank memory, likely with similar effects to increasing *q*; however, the effect of shifting the equilibrium may be stabilizing or destabilizing, depending on the underlying dynamics.

We focus on (local) equilibrium stability, which is only one facet of ecological stability [1, 5, 48]. Bilinsky and Hadeler [27] showed that dormancy can reduce the amplitude of predator-prey oscillations even when it is insufficient to qualitatively stabilize the equilibrium. Our geometric analysis provides new insight into this mechanism. Another important dimension of ecological stability is robustness to environmental forcing. The community matrix also influences how ecosystems respond to environmental noise [60], so we expect that dormancy might similarly dampen the response of food webs to stochastic forcing. And while our analysis suggests that dormancy is unlikely to stabilize competitive dynamics, seed banks can interact with environmental variability to facilitate coexistence of competing species via relative nonlinearity [61] or storage effects [62, 63, 22].

Overall, our analysis sheds new light on how and when we can expect dormancy to stabilize ecosystems. Dormancy suppresses oscillations associated with predator-prey dynamics (or, more generally, systems that lose stability through a Hopf bifurcation), and remains stabilizing in large, structured food webs, particularly when prey effects on predators are stronger than the reverse. Finally, we note that endogenous population fluctuations such as predator-prey oscillations can favor the evolution of dormancy as a bet-hedging strategy [64, 29, 26]. This suggests the potential for a negative feedback loop wherein fluctuations drive the evolution of dormancy, which in turn suppresses fluctuations – a mechanism by which food webs might self-organize to the edge of instability [26].

## Materials and Methods

### Deriving the ASD

We assume the underlying model dynamics have an equilibrium **x**^⋆^ *>* 0, meaning *f*_*i*_(**x**^⋆^) = 0 for all *i* = 1 … *n*. Then Eq. 1 has a corresponding equilibrium with 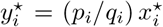. Letting 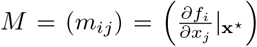 be the community (Jacobian) matrix for the underlying dynamics at **x**^⋆^, the system with dormancy (Eq. 1) has the community matrix

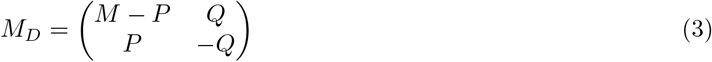

where *P* is a diagonal matrix with *P*_*ii*_ = *p*_*i*_ (and similarly for *Q*).

Under the simplifying assumption that *p*_*i*_ = *p* and *q*_*i*_ = *q* for all *i* (homogeneous rates), *P* and *Q* are scalar multiples of the identity matrix. If *λ* and **v** are an eigenvalue and corresponding eigenvector of *M*, then **v**_*D*_ = (**v**′, *k***v**′) is an eigenvector of *M*_*D*_ (with *k* an undetermined constant). Simplifying the eigenvalue equation *M*_*D*_**v**_*D*_ = *λ*_*D*_**v**_*D*_ and eliminating *k*, the eigenvalues *λ*_*D*_ satisfy the scalar equation

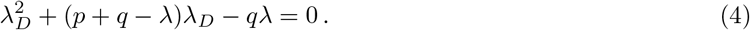

Eq. 4 relates each eigenvalue of *M* to two eigenvalues of *M*_*D*_, denoted by 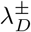, with 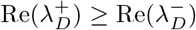. The set of possible *λ* with Re(*λ*) > 0 for which 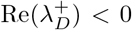 comprises the ASD (note that if Re(*λ*) < 0, then necessarily 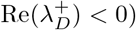.

The ASD boundary corresponds to solutions of Eq. 4 with Re(*λ*_*D*_) = 0. Hadeler and Lutscher [34] derived the parametric form

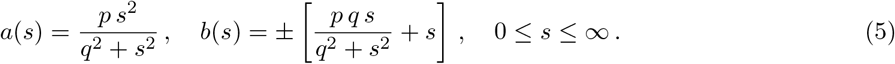

*λ*(*s*) = *a*(*s*) + *b*(*s*)*i* defines a family of solutions of Eq. 4 corresponding to purely imaginary 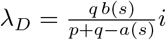. At *s* = 0, the ASD boundary passes through the origin. For *s* → ∞, we find the vertical asymptote *α* → *p, β* → ±∞. For more properties of the ASD, see [32, 33, 34, 35].

### Two-species models

We illustrate the effect of dormancy in two classical models, the symmetric Lotka-Volterra (LV) competition model and the MacArthur-Rosenzweig (MR) predator-prey model.

We define the LV model as

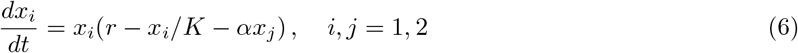

with intrinsic growth rate *r*, carrying capacity *K*, and interspecific competition strength *α. K* sets the strength of intraspecific competition. The LV model has an equilibrium with both species present at 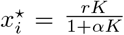. The eigenvalues of the community matrix at this equilibrium are *λ*_±_ = − (1*/K ± α*)*x*^⋆^. For *α* > 1*/K*, one eigenvalue is positive, indicating instability.

We define the MR model as

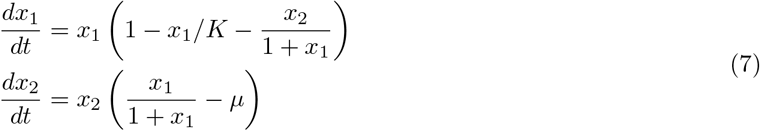

where species 1 is a prey that grows logistically in the absence of the predator (species 2). The prey’s carrying capacity in isolation is *K*, and the predator experiences fixed mortality *µ*. For *µ* < *K/*(*K* + 1), there is an equilibrium where 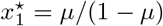 and 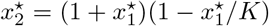. The eigenvalues of the community matrix can be expressed as 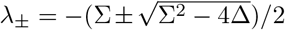, in terms of its trace 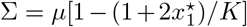 and determinant 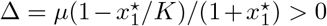 [38]. The coexistence equilibrium loses stability at 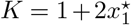, where Σ = 0. Because Δ > 0, the discriminant Σ^2^ − 4Δ is negative, indicating a Hopf bifurcation.

### Convergence of the seed bank to the time-averaged dynamics

The seed bank dynamics are governed by linear, first-order differential equations 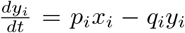, which admit the solutions

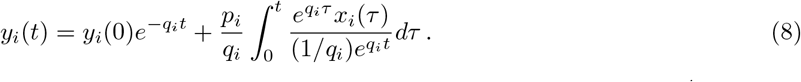

At long times (large *t*), the contribution of the initial conditions *y*_*i*_ (0) vanishes. Noting 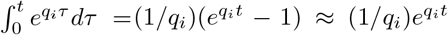, the seed bank state *y*_*i*_(*t*) converges to a (scaled) weighted average of the active dynamics: 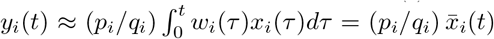 with weights 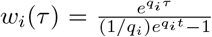 satisfying 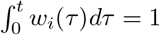. The active dynamics can then be approximated by Eq. 2 after an initial transient period.

### Random matrix models

To study the effect of dormancy in larger ecological networks, we model the community matrix *M* directly, allowing us to account for interaction strengths and structure while remaining agnostic to the specific dynamical model (e.g., the functional form of predator-prey interactions) [9, 7, 46, 56]. We consider three random matrix models and one structured food chain model (below). For random matrices, we assume complete connectance here for simplicity; allowing incomplete connectance has no qualitative effect on our results (SI Section 2A).

For unstructured mixed-sign interactions, we sample each element *m*_*ij*_ *iid* from a distribution with mean zero and variance *σ*^2^. For unstructured predator-prey interactions, we sample magnitudes |*m*_*ij*_| *iid* from a distribution with positive support, mean *µ*, and variance *σ*^2^, and then select one of *m*_*ij*_ and *m*_*ji*_ to have a negative sign at random (independently for each *i, j* pair). Note that the mean of *m*_*ij*_ is again zero. In both cases, we set *m*_*ii*_ = *S*, modeling self-regulation.

For any distribution of interaction strengths, as the number of species *n* becomes large, the eigenvalues of *M* in both unstructured cases are distributed uniformly over an ellipse in the complex plane defined by

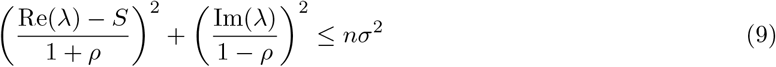

where *ρ* is the correlation between *m*_*ij*_ and *m*_*ji*_, a result known as the Elliptic Law [7, 41]. For the mixed-sign ensemble, *ρ* = 0, so the eigenvalues of *M* lie on disk centered at *S* with radius 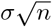 [9, 41]. For the predator-prey ensemble, 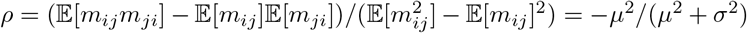. Thus, *ρ* < 0, and the eigenvalues lie in a vertically-stretched ellipse. In both cases, stability is controlled by eigenvalues with small imaginary parts with high probability.

To incorporate more realistic food web structure, we follow [46] and adopt the cascade model structure [42, 45], assuming that species are ordered in a trophic hierarchy from 1 to *n*. For each *i < j*, we sample *m*_*ij*_ from a distribution with negative support, mean *µ*_*U*_ *<* 0, and variance 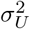, and *m*_*ij*_ from a distribution with positive support, mean *µ*_*L*_ *>* 0, and variance 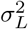 (independently). The negative elements *m*_*ij*_, in the upper-triangular part of *M*, model the effects of predators on prey; the positive elements *m*_*ji*_, in the lower-triangular part, model the effects of prey on predators. Again, we set *m*_*ii*_ = *S*.

Allesina et al. [46] showed that for *n* large, most eigenvalues fall into a circular “bulk” centered at *S*, with radius 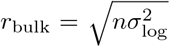, where 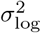 is the logarithmic mean 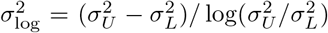. More generally, with connectance *C* < 1, the bulk is a vertically-stretched ellipse (as in Fig. 4). A smaller number of “outlier” eigenvalue pairs are located at

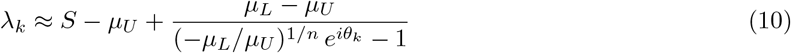

with *θ*_*k*_ = ± (2*k* − 1)*π/n* for *k* = 1, 2, …. The number of outliers depends on the size of the bulk. These eigenvalues lie on an arc in the complex plane passing through the bulk. Their behavior is more clearly revealed by the approximate form *λ*_*k*_ *≈ a*_*k*_ + *b*_*k*_*i* with

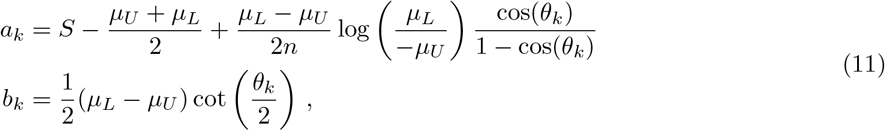

derived in SI Section 2B. The trigonometric factors in Eq. 11 are large when *θ*_*k*_ is small; thus, the first outlier(s) have the largest imaginary parts and are farthest from *S* (the center of the bulk). In the donor-controlled regime, |*µ*_*U*_ | *< µ*_*L*_, the logarithm in *a*_*k*_ is positive, so these large outliers fall to the right of the bulk. When |*µ*_*U*_| *> µ*_*L*_ (recipient-control), this orientation is reversed, and when |*µ*_*U*_| = *µ*_*L*_, the outliers lie on a vertical line.

For dormancy to have a stabilizing effect, it is necessary that *r*_bulk_ + *S <* 0 (the bulk is stabilized) and Re(*λ*_1_) *>* 0 (the rightmost outliers are destabilizing). In addition to |*µ*_*U*_ | *< µ*_*L*_ (donor-control), this demands that *r*_bulk_ is sufficiently small relative to Re(*λ*_1_). From Eq. 11, it is easy to see that changing |*µ*_*U*_| and *µ*_*L*_ in a fixed ratio scales the size of the outlier arc without changing its shape or orientation. Thus, the outliers control stability when mean interaction effects are sufficiently large relative to the logarithmic mean of their variances, which scales *r*_bulk_.

### Linear food chain model

As a minimal model of structured trophic interactions, we also study the eigenvalues of a simple linear food chain. We define the community matrix *M* where *m*_*i,i*+1_ = −*u* for 1 ≤ *i* ≤ *n* − 1 (predator effects on prey), *m*_*i,i*−1_ = *l* for 2 ≤ *i* ≤ *n* (prey effects on predators), *m*_*ii*_ = *S* for 2 ≤ *i* ≤ *n* − 1 (self effects). We distinguish *m*_11_ = *s*_*b*_ + *S* and *m*_*nn*_ = *s*_*a*_ + *S*, allowing enhanced self-regulation in the basal (*i* = 1) and apex species (*i* = *n*). All other *m*_*ij*_ are zero.

If *s*_*a*_ = *s*_*b*_ = 0, *M* is a tridiagonal Toeplitz matrix, with eigenvalues *λ* on a vertical line at Re(*λ*) = *S* [65]. Ecologically, we expect *s*_*a*_, *s*_*b*_ < 0, reflecting, for example, a carrying capacity in the basal species and “closure” effects in the top predator [10]. In SI Section 1A, we use a perturbation approach to approximate the eigenvalues when these terms are nonzero. We show that the line of eigenvalues bends, such that eigenvalues with large imaginary parts remain near Re(*λ*) = *S*, while those close to the real axis are deflected to the left. Thus, dormancy can be stabilizing if *S* > 0 and |*s*_*a*_ + *s*_*b*_| is sufficiently large relative to *S*. We expect *S* > 0 when trophic interactions are excitatory, e.g., when trophic levels are coupled by saturating functional responses. In particular, we show (SI Section 1B) that this *M* arises as the community matrix of an explicit dynamical model with Type II functional responses and high trophic transfer efficiency.

## Acknowledgements

We thank A. Skwara and S. Allesina for helpful comments and discussion. This research received support through Schmidt Sciences LLC.

