## Supplemental Information for "Dormancy stabilizes structured food webs"

### Supplementary information for:
### Dormancy stabilizes structured food webs

Zachary R. Miller[1]*, David A. Vasseur[2], Pincelli M. Hull[1]
[1]Department of Earth & Planetary Sciences, Yale University, New Haven, CT 06511
[2]Department of Ecology & Evolutionary Biology, Yale University, New Haven, CT 06511
*

May 28, 2026

**4 Stabilization of intransitive competition** **13**

**5 Additional parameter and simulation details** **18**

### 1 Linear food chain model

#### 1A Approximating eigenvalues of the community matrix

We are interested in the eigenvalues of a community matrix corresponding to a linear food chain of length $n$, which we model as

$$M = \begin{pmatrix} S+s_b & -u & & & \\ l & S & -u & & 0 \\ & l & \ddots & \ddots & \\ 0 & & \ddots & S & -u \\ & & & l & S+s_a \end{pmatrix} \tag{1}$$

with $u, l > 0$. We leave the signs of $S, s_a$, and $s_b$ arbitrary, although we will be most interested in cases where $S > 0$ and $s_a + s_b < 0$.

As explained in the Main Text (Materials and Methods), $-u$ models the effect of predators on prey, $l$ models the effects of prey on predators, and $S$ models self-interaction effects. $s_b$ and $s_a$ allow modified self-effects in the basal and apex species, respectively. It is natural to assume that there is enhanced self-regulation in the basal species $s_b < 0$, reflecting a carrying capacity in the absence of higher trophic levels, and also in the top predator $s_a < 0$, reflecting unmodeled density-dependent sources of mortality [1].

While conditions for food chain stability and persistence (e.g., permanence) have been well-studied using more powerful approaches than local stability analysis [2, 3, 1], to the best of our knowledge, the set of eigenvalues, or *spectrum*, of matrices like (1) has not been characterized in the ecological literature. Matrices of this form have received attention in other fields, leading to a substantial body of results on their eigenvalues and eigenvectors [4, 5, 6, 7]. While these include explicit formulas for the eigenvalues in special cases (e.g., $s_a s_b = -ul$), we are not aware of any explicit formulas for the general case that provide clear insight into the geometry of the spectrum for finite $n$. Thus, we develop a novel approximation for these eigenvalues.

In the degenerate case where $s_a, s_b = 0$, $M$ becomes a tridiagonal Toeplitz matrix. The eigenvalues of this type of matrix are well-known, and can be found exactly by solving a system of linear difference equations. These eigenvalues are

$$\lambda_k = S - 2\sqrt{ul}\cos\left(\frac{k\pi}{n+1}\right) i\,, \quad k = 1, 2, \ldots n. \tag{2}$$

which lie on a vertical line in the complex plan with real part $S$. Some of these eigenvalues have very small negative parts (e.g., consider $k \approx (n+1)/2$, in which case $\cos(\pi/2) = 0$), indicating that dormancy cannot stabilize this spectrum.

To characterize the eigenvalues of $M$ with at least one of $s_a, s_b$ nonzero, we approximate them as a perturbation of the tridiagonal Toeplitz case.

We begin by reducing the number of parameters. Consider the matrix $M' = (1/\sqrt{ul})(M - SI)$, where $I$ is the identity matrix. The eigenvalues of $M$ and $M'$ are related by $\lambda' = (1/\sqrt{ul})(\lambda - S)$. Defining $x = s_b/\sqrt{ul}$, $y = s_a/\sqrt{ul}$, and $z = \sqrt{u/l}$, we have

$$M' = \begin{pmatrix} x & -z & & & \\ 1/z & 0 & -z & & \\ & 1/z & \ddots & \ddots & \\ & & \ddots & 0 & -z \\ & & & 1/z & y \end{pmatrix}. \tag{3}$$

We now write the characteristic polynomial $p_n^{xy}(\lambda) = \det(M' - \lambda I)$ in a recursive form. Expanding the determinant along the first row of $M'$, we have

$$p_n^{xy} = (x - \lambda) p_{n-1}^{0y} + p_{n-2}^{0y}\,. \tag{4}$$

Proceeding recursively,

$$p_k^{0y} = -\lambda p_{k-1}^{0y} + p_{k-2}^{0y} \tag{5}$$

with initial conditions $p_0^{0y} = 1$ and $p_1^{0y} = y - \lambda$. The linear difference equation (5) admits solutions of the form $p_k^{0y} = c_+ t_+^k + c_- t_-^k$ with

$$t_\pm = \frac{-\lambda \pm \sqrt{\lambda^2+4}}{2}\,, \tag{6}$$

obtained by solving the associated characteristic equation. Using the initial conditions and the fact that $-\lambda = t_+ + t_-$, we find

$$c_\pm = \frac{y + t_\pm}{t_+ - t_-}. \tag{7}$$

Finally, substituting the solution for $p_k^{0y}$ into Eq. 4, we have the characteristic polynomial

$$p_n^{xy}(\lambda) = \frac{(x-\lambda)(t_+^n - t_-^n) + (xy - y\lambda + 1)(t_+^{n-1} - t_-^{n-1}) + y(t_+^{n-2} - t_-^{n-2})}{t_+ - t_-}\,. \tag{8}$$

It is clear already from this expression that the qualitative behavior of the eigenvalues will have no dependence on $z$ (the ratio of $u$ to $l$). However, $x, y \neq 0$ cause a non-trivial deviation from the Toeplitz case.

We assume that $t_+ \neq t_-$ and seek eigenvalues $\lambda$ satisfying $p_n^{xy}(\lambda) = 0$. To express $p_n^{xy}$ in a more convenient form, note that $t_+ t_- = -1$. Denoting $t = t_+$, we have $t_- = -1/t$ and $\lambda = 1/t - t$. Thus, we can write

$$p_n^{xy}(t) = t^{n+1} + (x+y)t^n + xy\, t^{n-1} \pm xy\, t^{-(n-1)} \mp (x+y)t^{-n} \pm t^{-(n+1)} \tag{9}$$

for $n$ even and odd, respectively.

We are unable to solve this polynomial equation in general. Instead, we attempt to approximate solutions of $p_n^{xy}(t) = 0$ using the ansatz that, for $|x|, |y| < 1$, these roots lie near those for the Toeplitz case, where $x, y = 0$. Defining $\theta_k = k\pi/(2n+2)$, these solutions are

$$t_k^* = \exp(i\theta_k)\,, \quad k = -(n-1), -(n-3), \ldots, n-1\,. \tag{10}$$

We Taylor-expand $p_n^{xy}(t)$ to first-order around $t_k^*$:

$$\begin{aligned} p_{n,k}^{xy}(t) \approx{}& \exp((n+1)\,i\,\theta_k) \pm \exp(-(n+1)\,i\,\theta_k) + (n+1)\exp(-i\theta_k)\Big(\exp((n+1)\,i\,\theta_k) \mp \exp(-(n+1)\,i\,\theta_k)\Big)(t - t_k^*) + \\ & (x+y)\Big[\exp(n\,i\,\theta_k) \mp \exp(-n\,i\,\theta_k) + n\exp(-i\theta_k)\Big(\exp(n\,i\,\theta_k) \pm \exp(-n\,i\,\theta_k)\Big)(t - t_k^*)\Big] + \\ & xy\Big[\exp((n-1)\,i\,\theta_k) \pm \exp(-(n-1)\,i\,\theta_k) + (n-1)\exp(-i\theta_k)\Big(\exp((n-1)\,i\,\theta_k) \mp \exp(-(n-1)\,i\,\theta_k)\Big)(t - t_k^*)\Big] \end{aligned} \tag{11}$$

Recognizing trigonometric functions and solving for $t$ with $p_{n,k}^{xy}(t) = 0$, we eventually find the approximate roots

$$\tilde{t}_k = \begin{cases} t_k^*\left(1 - \frac{(x+y)\,i\,\sin(n\theta_k) + xy\,\cos((n-1)\theta_k)}{(n+1)\,i\,\sin((n+1)\theta_k) + (x+y)n\cos(n\theta_k) + xy\,(n-1)\,i\,\sin((n-1)\theta_k)}\right)\,, & n \text{ even} \\ t_k^*\left(1 - \frac{(x+y)\,i\,\cos(n\theta_k) + xy\,\sin((n-1)\theta_k)}{(n+1)\,i\,\cos((n+1)\theta_k) + (x+y)n\sin(n\theta_k) + xy\,(n-1)\,i\,\cos((n-1)\theta_k)}\right)\,, & n \text{ odd} \end{cases} \tag{12}$$

From Eq. 12, we can calculate approximate eigenvalues $\tilde{\lambda}_k$ using $\tilde{\lambda}_k = 1/\tilde{t}_k - \tilde{t}_k$. Shifting and rescaling appropriately, we obtain approximate eigenvalues for the original matrix $M$. Fig. 1 shows that this yields a good approximation across a range of $n$ when $x$ and $y$ are sufficiently small.

To better understand the qualitative behavior of the eigenvalues, we make a further approximation: For moderately large $n$, we can very accurately approximate the trigonometric terms around $(n+1)/(n+1) = 1$; for example,

$$\begin{aligned} \cos(n\,\theta_k) &= \cos\left(\frac{n}{n+1}\frac{k\pi}{2}\right) \approx 0 - \frac{k\pi}{2}\sin\left(\frac{k\pi}{2}\right)\frac{-1}{n+1} = \pm\theta_k\,, \quad n \text{ even } (k \text{ odd}) \\ \sin(n\,\theta_k) &= \sin\left(\frac{n}{n+1}\frac{k\pi}{2}\right) \approx 0 + \frac{k\pi}{2}\cos\left(\frac{k\pi}{2}\right)\frac{-1}{n+1} = \mp\theta_k\,, \quad n \text{ odd } (k \text{ even}) \end{aligned} \tag{13}$$

and so on. This yields

$$\tilde{t}_k \approx t_k^*\left(1 - \frac{(x+y) - 2xy\,\theta_k\,i}{(n+1) + (n-1)xy - n(x+y)\,\theta_k\,i}\right) \approx t_k^*\left(1 - \frac{(x+y) - 2xy\,\theta_k\,i}{n(1 + xy - (x+y)\,\theta_k\,i)}\right) \tag{14}$$

for all $n$ (even or odd).

From Eq. 14, some qualitative features of the spectrum become apparent. First, note that our approximation is of the form $\tilde{t}_k \approx t_k^*(1-\delta/n)$, i.e., a small proportional perturbation. Some arithmetic shows that $\mathrm{Re}(\delta) \propto (x+y)(1+xy(1+2\theta_k^2))$. If we assume that $x, y < 0$, then clearly $\mathrm{Re}(\delta) < 0$ and thus $\mathrm{Re}(1-\delta/n) > 1$. The values $t_k^*$ lie on a semicircle of radius one in the right-half of the complex plane – the effect of the small perturbation is to "inflate" this semicircle. All $\tilde{t}_k$ lie near $t_k^*$, but with slightly larger modulus. Finally, we consider how this perturbation is propagated through the transformation

$$\lambda = 1/t - t = \left(\frac{1}{|t|^2} - 1\right)\mathrm{Re}(t) + \left(\frac{1}{|t|^2} + 1\right)\mathrm{Im}(t)\, i \, . \tag{15}$$

We know that $|\tilde{t}_k| > 1$ and $\mathrm{Re}(\tilde{t}_k) > 0$. Thus, $\mathrm{Re}(\lambda_k) < 0$. Geometrically, this transformation reflects the (approximate) semicircle formed by the $\tilde{t}_k$ across the imaginary axis, and stretches it vertically (because we have $1/|\tilde{t}_k| - 1$ small and $1/|\tilde{t}_k| + 1 > 1$). Thus, we find a crescent-shaped curve opening to the right, as seen in Fig. 3 (Main Text) and Fig. 1.

To recover approximate eigenvalues of $M$ from these $\tilde{\lambda}_k$, we multiply by $\sqrt{ul}$ and shift by $+S$. The rightmost $\tilde{\lambda}_k$ lie near the imaginary line; thus, $S > 0$ will push the rightmost eigenvalues into the right half-plane. If $S$ is not too large, then the eigenvalues with small imaginary parts remain in the left half-plane, creating the conditions for dormancy to have a stabilizing effect. In particular, for $n$ odd, there is a real eigenvalue corresponding to $k = 0$; from our approximation, this eigenvalue is approximately (and strictly less than) $(x+y)/[n(1+xy)]$. Dormancy is therefore most likely to have a stabilizing effect when $0 < S < |s_a + s_b|/[n(1 + s_a s_b/ul)]$ with $s_a + s_b < 0$.

#### 1B Derivation from a dynamical model with saturating functional responses

Here, we show that community matrices of the form of Eq. 1 can arise from an (idealized) dynamical food chain model. Consider the quite general food chain model

$$\begin{aligned}
\frac{dx_1}{dt} &= x_1\Big(r - g_1(x_1) - f(x_1)x_2\Big) \\
\frac{dx_i}{dt} &= x_i\Big(\epsilon f(x_{i-1})x_{i-1} - f(x_i)x_{i+1} - d_i\Big)\,, \quad i = 2,\dots,n-1 \\
\frac{dx_n}{dt} &= x_n\Big(\epsilon f(x_{n-1})x_{n-1} - g_n(x_n) - d_n\Big)\,,
\end{aligned} \tag{16}$$

where species 1 is the basal species and species $n$ is the top predator. All parameters and functions are non-negative. We are assuming that the functional response, $f$, coupling trophic levels is identical at each level of the food chain. We will also assume that $g_1' \geq 0$ and $g_n' \geq 0$, modeling negative density-dependence. Let $x_i^\star$ be an equilibrium of Eq. 16. The community matrix at equilibrium is given by

$$\begin{aligned}
m_{11} &= -x_1^\star g_1'(x_1^\star) - f'(x_1^\star)x_1^\star x_2^\star \\
m_{i,i+1} &= -x_i^\star f(x_i^\star)\,, && i = 1,\dots,n-1 \\
m_{ii} &= -f'(x_i^\star)x_i^\star x_{i+1}^\star\,, && i = 2,\dots,n-1 \\
m_{i,i-1} &= \epsilon x_i^\star(f(x_{i-1}^\star) + f'(x_{i-1}^\star)x_{i-1}^\star)\,, && i = 2,\dots,n \\
m_{nn} &= -x_n^\star g_n'(x_n^\star)
\end{aligned} \tag{17}$$

with all other $m_{ij} = 0$. Clearly, this community matrix has the same tridiagonal structure as Eq. 1. Additionally, if we assume that $f' < 0$ (saturating functional response), then $m_{ii} > 0$, motivating the case $S > 0$. We also have $m_{i,i+1} < 0$, while the sign of $m_{i,i-1}$ depends on the relative values of $f$ and $f'$ at equilibrium.

If we make the idealized assumptions that $\epsilon \approx 1$ (very high trophic transfer efficiency) and $d_i \approx 0$ for $i = 2,\dots,n-1$ (losses to predation are much larger than other sources of mortality), then $f(x_{i-1}^\star)x_{i-1}^\star \approx f(x_i^\star)x_{i+1}^\star$ at equilibrium, which may be satisfied by $x_i \approx x_j$ for all $i, j$. In this case, where $m_{ii} \approx m_{jj}, m_{i,i-1} \approx m_{j,j-1}$, and $m_{i,i+1} \approx m_{j,j+1}$ for all $i, j$, we may view Eq. 1 as an approximation of the community matrix.

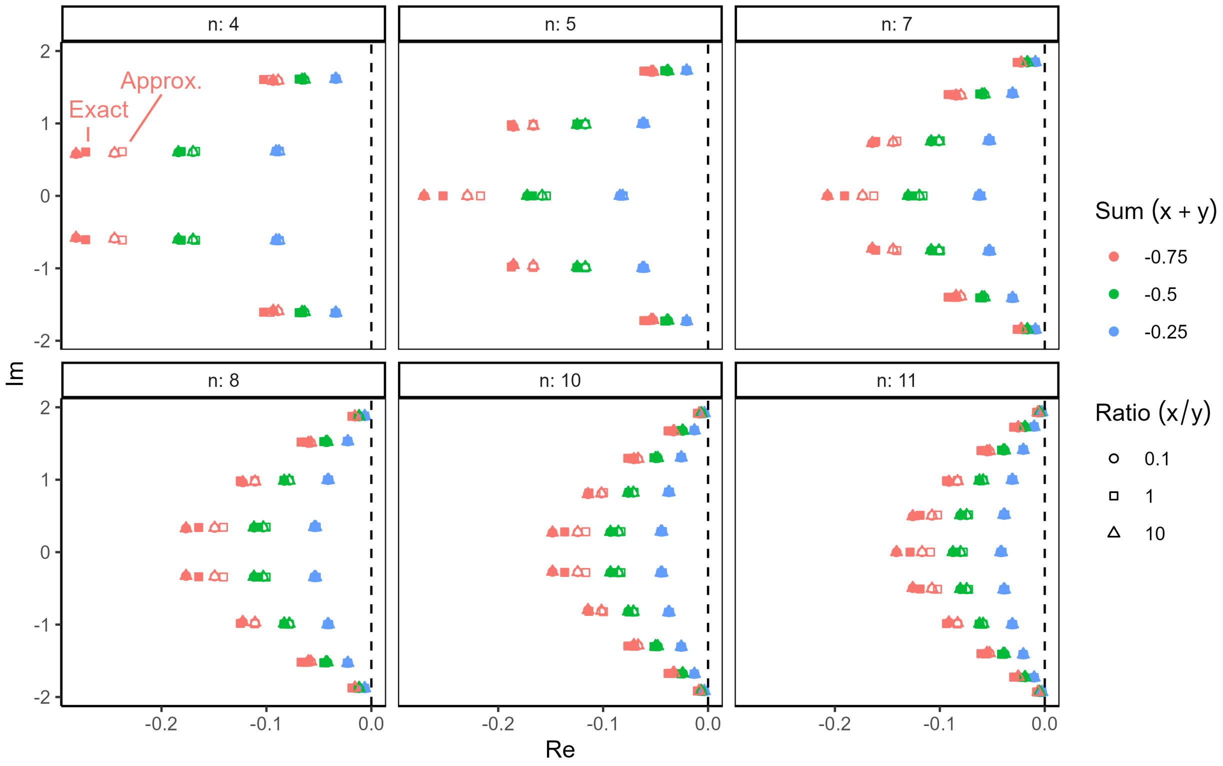

Figure 1: Comparison of exact and approximate eigenvalues of the simplified food chain community matrix (Eq. 3). We plot exact eigenvalues, calculated using numerical eigendecomposition in R (filled shapes), alongside the approximation based on Eq. 12 (open shapes) for different food chain lengths $n$ (panels), and sums (colors) and ratios (shapes) of $x$ and $y$. This parameterization highlights that the eigenvalues depend primarily on the sum $x + y$, and only weakly on the ratio $x/y$. In all cases, the qualitative behavior of the spectrum is the same: eigenvalues lie on a crescent-shaped curve, with the rightmost eigenvalues having large imaginary parts. For increasing $|x + y|$, the deviation between our approximation and the exact eigenvalues increases, but we note that these deviations remain consistent with the qualitative behavior predicted by our approximation theory. In fact we observe greater curvature in the exact spectrum, indicating more potential for dormancy to stabilize the corresponding dynamics. Eigenvalues of matrix (1) can be obtained from these values by appropriate shifting and scaling, as described in the text.

Let us consider one specific example where we recover Eq. 1 exactly. We assume a Type II functional response, i.e., $f(x) = \alpha/(h+x)$. We again assume that $\epsilon \approx 1$, $d_i \approx 0$ for $i = 2, \ldots, n-1$, and additionally $x_1^\star \approx x_2^\star \approx \cdots \approx x_n^\star = x$ (equal biomass at each trophic level). The last assumption entails that $r = \alpha x/(h+x) + g_1(x)$ and $d_n = \alpha x/(h+x) - g_n(x)$, which can always be achieved through an appropriate choice of $r$ and $d_n$ (noting that we must have $\alpha x/(h+x)/g_n(x)$ in order for $d_n > 0$). Under this parameterization, we have the community matrix elements

$$\begin{aligned}
m_{11} &= -x g_1'(x) + \frac{\alpha x^2}{(h+x)^2} \equiv s_b + S \\
m_{i,i+1} &= -\frac{\alpha x}{h+x} \equiv -u\,, && i = 1, \ldots, n-1 \\
m_{ii} &= \frac{\alpha x^2}{(h+x)^2} \equiv S\,, && i = 2, \ldots, n-1 \\
m_{i,i-1} &= \frac{\alpha h x}{(h+x)^2} \equiv l\,, && i = 2, \ldots, n \\
m_{nn} &= -x g_n'(x) \equiv s_a + S
\end{aligned} \tag{18}$$

which have the same signs and structure as Eq. 1. Notice that this implies $s_a < 0$ even if $g_n = 0$.

### 2 Random matrix models: Additional details

#### 2A Effect of connectance

In the Main Text, we assume complete connectance in our random networks for simplicity of presentation. Here we detail the effect of allowing incomplete connectance, as is more realistic for most ecological networks (especially trophic networks). This change produces quantitative corrections to the spectra, but has no qualitative effect on our results or conclusions.

In all three of our random matrix ensembles, we can model incomplete connectance by choosing to set $m_{ij} = 0$ and $m_{ji} = 0$ with probability $1 - C$ for each pair $i, j$ independently. For the two unstructured cases, which have eigenvalues following the Elliptic Law (Eq. 9, Main Text), incomplete connectance simply has the effect of scaling the variance $\sigma^2$ by $C$. Thus, the eigenvalues are again distributed over an ellipse, now given by

$$\left(\frac{\mathrm{Re}(\lambda) - S}{1+\rho}\right)^2 + \left(\frac{\mathrm{Im}(\lambda)}{1-\rho}\right)^2 \leq nC\sigma^2\,. \tag{19}$$

For the cascade model matrices, the effect of connectance is slightly more involved [8]. Calculating the means, variances, and correlation of upper and lower triangular elements of $M$ after accounting for incomplete connectance $C$, we find

$$\begin{aligned}
\hat{\mu}_U &= C\mu_U \\
\hat{\mu}_L &= C\mu_L \\
\hat{\sigma}_U^2 &= C(\sigma_U^2 + \mu_U^2(1-C)) \\
\hat{\sigma}_L^2 &= C(\sigma_L^2 + \mu_L^2(1-C)) \\
\hat{\rho} &= -\frac{C(1-C)\mu_U\mu_L}{\hat{\sigma}_U\hat{\sigma}_L}
\end{aligned} \tag{20}$$

respectively, where the "unhatted" values refer to the means and variances of the underlying distributions and the "hatted" values account for incomplete connectance. We now find a nonzero correlation between values $m_{ij}$ and $m_{ji}$. In this case, the bulk of eigenvalues takes the shape of an ellipse, rather than a circle. From [8], the horizontal and vertical axes of the ellipse are given by

$$r_h = \frac{n\hat{\sigma}_{\log}^2 + \hat{\rho}\hat{\sigma}_U\hat{\sigma}_L(n-1)}{\sqrt{n\hat{\sigma}_{\log}^2}} \qquad r_v = \frac{n\hat{\sigma}_{\log}^2 - \hat{\rho}\hat{\sigma}_U\hat{\sigma}_L(n-1)}{\sqrt{n\hat{\sigma}_{\log}^2}} \tag{21}$$

where $\hat{\sigma}^2_{\log}$ is the logarithmic mean $\hat{\sigma}^2_{\log} = (\hat{\sigma}^2_U - \hat{\sigma}^2_L)/\log(\hat{\sigma}^2_U/\hat{\sigma}^2_L)$. We always have $\hat{\rho} \leq 0$, so incomplete connectance tends to compress the bulk in the horizontal direction (see Fig. 4, Main Text). The outliers are found as before using Eq. 10 in the Main Text, but substituting in the hatted means. Note that while the bulk now has some dependence on the means $\mu_U$ and $\mu_L$ (which appear in the formulae for $\hat{\sigma}^2_U, \hat{\sigma}^2_L$ and $\hat{\rho}$ under incomplete connectance), the outliers remain unaffected by the interaction variances.

#### 2B Approximating outliers of the random cascade model

Allesina et al. [8] showed that the outlier eigenvalues of random cascade model matrices are located approximately at

$$\lambda_k \approx S - \mu_U + \frac{\mu_L - \mu_U}{(-\mu_L/\mu_U)^{1/n}\, e^{i\theta_k} - 1} \tag{22}$$

with $\theta_k = \pm(2k-1)\pi/n$ for $k = 1, 2, \ldots$. The number of outliers (maximum value of $k$) depends on the size of the bulk.

Here, we derive an approximation for these outliers that more clearly reveals their geometric behavior in the complex plane. Allesina et al. [8] also showed that the real and imaginary parts of these eigenvalues can be written separately as

$$\mathrm{Re}(\lambda_k) = S - \mu_U \left(1 + \frac{(r+1)(r^{1/n}\cos\theta_k - 1)}{1 + r^{2/n} - 2r^{1/n}\cos\theta_k}\right) \qquad \mathrm{Im}(\lambda_k) = \mu_U \left(\frac{(r+1)r^{1/n}\sin\theta_k}{1 + r^{2/n} - 2r^{1/n}\cos\theta_k}\right) \tag{23}$$

where $r = -\mu_L/\mu_U$. For large $n$, we have the very good approximation $r^{1/S} \approx 1 + \log(r)/n$, which follows from a first-order Taylor expansion of $r^{1/S}$ written as $\exp(\log(r)/n)$. Using this approximation, and neglecting small (i.e., $\log(r)/n$) terms in the denominators, we have

$$\mathrm{Re}(\lambda_k) \approx S - \mu_U \left(1 + \frac{(r+1)\Big((1 + \log(r)/n)\cos\theta_k - 1\Big)}{2(1 - \cos\theta_k)}\right) \qquad \mathrm{Im}(\lambda_k) \approx \mu_U \left(\frac{(r+1)(1 + \log(r)/n)\sin\theta_k}{2(1 - \cos\theta_k)}\right) \tag{24}$$

Some re-arranging and substituting $r = -\mu_L/\mu_U$ yields

$$\mathrm{Re}(\lambda_k) \approx S - \frac{\mu_U + \mu_L}{2} + \frac{\mu_L - \mu_U}{2n} \log\left(\frac{\mu_L}{-\mu_U}\right) \frac{\cos(\theta_k)}{1 - \cos(\theta_k)}\,. \tag{25}$$

Clearly, the real part of the eigenvalues depends on $k$ only through the last term. As discussed in the Main Text, for the first few values of $k$, $\theta_k \approx 0$, and thus the ratio $\frac{\cos(\theta_k)}{1-\cos(\theta_k)}$ is very large. The sign of this term is controlled by $\log\left(\frac{\mu_L}{-\mu_U}\right)$. If $\mu_L > -\mu_U$ (donor-controlled regime), then the logarithm is positive. Putting these facts together, the first few outliers (first few $k$) fall to the right of the rest only in the donor-controlled regime. These eigenvalues also have large imaginary parts, as we can see by using half-angle identities to recognize the cotangent and writing

$$\mathrm{Im}(\lambda_k) \approx \frac{1}{2}(\mu_L - \mu_U)\cot\left(\frac{\theta_k}{2}\right)\,. \tag{26}$$

For small $\theta_k$ the cotangent is very large in magnitude (with the same sign as $\theta_k$).

### 3 Rate heterogeneity

#### 3A No stabilization of symmetrizable community matrices

With heterogeneous rates $p_i$ and $q_i$, it is difficult to characterize the stability of equilibria of Eq. 1 (Main Text) in general. As discussed by Hadeler [9], this problem is closely related to the well-studied Turing (in)stability problem and the problem of characterizing strong matrix stability; thus, we would be surprised to find a straightforward, general characterization.

However, we can provide a partial answer. We prove that if the underlying community matrix $M$ is *symmetrizable* and unstable, then no choice of (in)activation rates can stabilize the system. A symmetrizable matrix is one that can be written as $M = D_1 A D_2$, where $A$ is a real, symmetric matrix and $D_1, D_2$ are real, positive diagonal matrices. This type of matrix arises naturally in Lotka-Volterra models with symmetric interactions because the Jacobian matrix is multiplied by species' abundances at equilibrium [10]. As a particular example, symmetrizable community matrices arise when interactions reflect implicit resource competition with substitutable biotic resources [11, 12].

Our strategy of proof is to show that a symmetrizable system with heterogeneous rates is stable if and only if an associated system with homogeneous rates is stable. Then, we are able to use the ASD to classify stability.

If $M$ is symmetrizable, then its eigenvalues are purely real because $M$ is similar to a symmetric matrix: we have $BMB^{-1} = D_1^{1/2} D_2^{1/2} A D_2^{1/2} D_1^{1/2}$ with $B = D_1^{-1/2} D_2^{1/2}$. Consider the community matrix for the system with dormancy:

$$M_D = \begin{pmatrix} D_1 A D_2 - P & Q \\ P & -Q \end{pmatrix} . \tag{27}$$

$M_D$ is also similar to a symmetric matrix, as we can see:

$$M'_D = \begin{pmatrix} B & 0 \\ 0 & B\, Q^{1/2} P^{-1/2} \end{pmatrix} M_D \begin{pmatrix} B^{-1} & 0 \\ 0 & B^{-1}\, Q^{-1/2} P^{1/2} \end{pmatrix} = \begin{pmatrix} D_1^{1/2} D_2^{1/2} A D_2^{1/2} D_1^{1/2} - P & P^{1/2} Q^{1/2} \\ P^{1/2} Q^{1/2} & -Q \end{pmatrix} . \tag{28}$$

Now we pre- and post-multiply $M'_D$ by another block diagonal matrix to obtain

$$M''_D = \begin{pmatrix} P^{-1/2} & 0 \\ 0 & Q^{-1/2} \end{pmatrix} M'_D \begin{pmatrix} P^{-1/2} & 0 \\ 0 & Q^{-1/2} \end{pmatrix} = \begin{pmatrix} P^{-1/2} D_1^{1/2} D_2^{1/2} A D_2^{1/2} D_1^{1/2} P^{-1/2} - I & I \\ I & -I \end{pmatrix} . \tag{29}$$

Because $M'_D$ is symmetric, $M'_D$ and $M''_D$ have the same number of positive eigenvalues by Sylvester's law of inertia. $M_D$ and $M'_D$ have precisely the same eigenvalues by similarity. Thus, if $M''_D$ is unstable then so is $M_D$.

$M''_D$ has the same form as our eigenvalue problem with homogeneous rates (see Materials and Methods in the Main Text) with $p = q = 1$. Consequently, if the symmetric matrix $A'' = P^{-1/2} D_1^{1/2} D_2^{1/2} A D_2^{1/2} D_1^{1/2} P^{-1/2}$ is unstable, then it must have at least one real positive eigenvalue, and we know that this eigenvalue must fall outside of the ASD. $A''$ is congruent to $A' = D_1^{1/2} D_2^{1/2} A D_2^{1/2} D_1^{1/2}$ (i.e., pre- and post-multiplied by an invertible matrix and its transpose, which in this case are identical), so they have the same number of positive eigenvalues, again by Sylvester's law of inertia. But $A'$ and $M$ are similar, as we showed above, so we conclude that if $M$ has a positive real eigenvalue, then so does $A''$, and thus $M''_D$ is unstable.

It is important to note that this proof assumes $p_i, q_i > 0$ for all $i$. However, our argument applies with arbitrarily small (positive) rates; by continuity, we expect that allowing some rates exactly equal to zero will not change the qualitative stability behavior.

We also note that our proof implies that for $n = 2$, only predator-prey $(+, -)$ interactions can be stabilized by dormancy. That is, only community matrices where $m_{12} m_{21} < 0$. This is because any $2 \times 2$ matrix with with $m_{12} m_{21} > 0$ is symmetrizable: we can write

$$M = \begin{pmatrix} a & b \\ c & d \end{pmatrix} = \begin{pmatrix} \sqrt{b/c} & 0 \\ 0 & \sqrt{c/b} \end{pmatrix} \begin{pmatrix} a\sqrt{c/b} & \sqrt{bc} \\ \sqrt{bc} & d\sqrt{b/c} \end{pmatrix} \tag{30}$$

if and only if $b$ and $c$ have the same sign.

#### 3B Supplementary simulation results

In this section, we report additional simulation results for the random cascade model with heterogeneous dormancy rates.

Fig. 2 shows the threshold behavior of stabilization in the donor-controlled (D) regime. The likelihood that dormancy stabilizes the model food webs is strongly predicted by the product $pF$, where $F$ is the fraction of species in the community that have the capacity to go dormant and $p$ is the inactivation rate for those that do.

Clearly, specific attributes of the set of species with dormancy can only have a substantial effect on the probability of stability in a narrow range around $pF = \text{Re}_{\text{max}}$. To explore these effects, we quantified several characteristics of the set of species with dormancy across all simulations for $p - F$ combinations where the overall probability of stabilization was more than 25% but less than 75%. Specifically, for each species with dormancy, we calculated its degree in the interaction network (the number of interaction partners, including predators and prey), the overall strength of each its interactions (quantified as $\sum_j |m_{ij}m_{ji}|$), and its trophic rank. To summarize the *set* of species with dormancy, we considered both the mean and maximum value of these attributes across all species in the set. For trophic rank, we also considered the minimum. These potential predictors were chosen to assess several intuitive hypothesis about the effect of dormancy – for example, that dormancy is most stabilizing when it occurs in highly connected species, strongly interacting species, top predators, basal species, etc.

In Fig. 3, we plot the distribution of these set-level attributes conditionally on whether dormancy stabilized the network or not. We see a separation in degree and interaction strength (both mean and max) – these quantities are higher on average in cases where dormancy is stabilizing – but not for trophic rank. This suggests that dormancy may be more stabilizing when the species that go dormant are more connected and interact more strongly with other species in the food web (whether as prey or predator).

To further explore these hypotheses, we examined the relationship between these attributes and the probability of stabilization in three different $p - F$ combinations along the curve $pF = \text{Re}_{\text{max}}$ (i.e., three pixels along the green dashed curve in Fig. 4, Main Text). We visualize this relationship in Fig. 4 by fitting a logistic regression with stable vs. unstable outcomes and each subset attribute as a predictor (individually). We see that the probability of stability increases sharply as a function of degree (mean or max) and interaction strength (mean or max), but not trophic rank.

Finally, we examined the predictive power of these subset attributes for individual food webs (i.e., realizations of the community matrix). Fig. 5 shows stability vs. subset attributes for three randomly chosen food webs within each of the three $p - F$ combinations shown in Fig. 4. Each point represents one randomly chosen subset of species (8, 12, or 16 species) with dormancy for the same underlying community matrix (25 for each web). As in Fig. 4, we fit a logistic regression to visualize the effect of the predictors (we only show mean degree, interaction strength, and trophic rank for clearer visualization). Here, fixing $p$, $F$, and the web identity, we see that degree and interaction strength are excellent predictors of stability. In some cases (e.g., $F : 0.32\,(1, 3)$ and $F : 0.64\,(1)$), we find complete separation, so that all subsets below a threshold value fail to stabilize the network, while all of those above produce stabilization. While these threshold points vary by web, this suggests that the stabilizing effect of dormancy in these cases may be highly predictable from the degree or average interaction strength of the species with dormancy. Once again at this granular level, we find that trophic rank has no consistent predictive power.

In contrast to the donor-controlled (D) regime, in the recipient-controlled (R) and balanced interaction (B) regimes, the probability of stability is never high. As discussed in the Main Text, this is a result of the different outlier eigenvalue geometry in these three regimes. In Fig. 6, we show that stabilization in the R and B regimes only occurs in a small number of food webs for which the destabilizing eigenvalues (those with positive real parts) all have relatively large imaginary parts. This variation between food webs occurs due to chance fluctuations in the "bulk" of eigenvalues.

We also conducted two additional sets of simulations to explore the effect of varying some assumptions made in the Main Text.

In Fig. 7, we repeat the simulations shown in Fig. 4, but with $q = 2$. We find qualitatively similar results. For this substantially larger value of $q$, the asymptotic phase of the ASD is pushed farther from the real line. Thus, stabilization by dormancy becomes less likely at each value of $p - F$. In the homogeneous case ($F = 1$), stabilization of donor-controlled webs occurs around $p \approx 0.22$. As in the Main Text, we find that stabilization becomes highly likely once the product $pF$ exceeds this (larger) value. Stabilization in the B and R regimes is very unlikely.

In Fig. 8, we repeat these simulations once again, but relaxing the assumption that species either do not have the capacity for dormancy or go dormant with the same rates $p$ and $q$ as all others. Instead, we

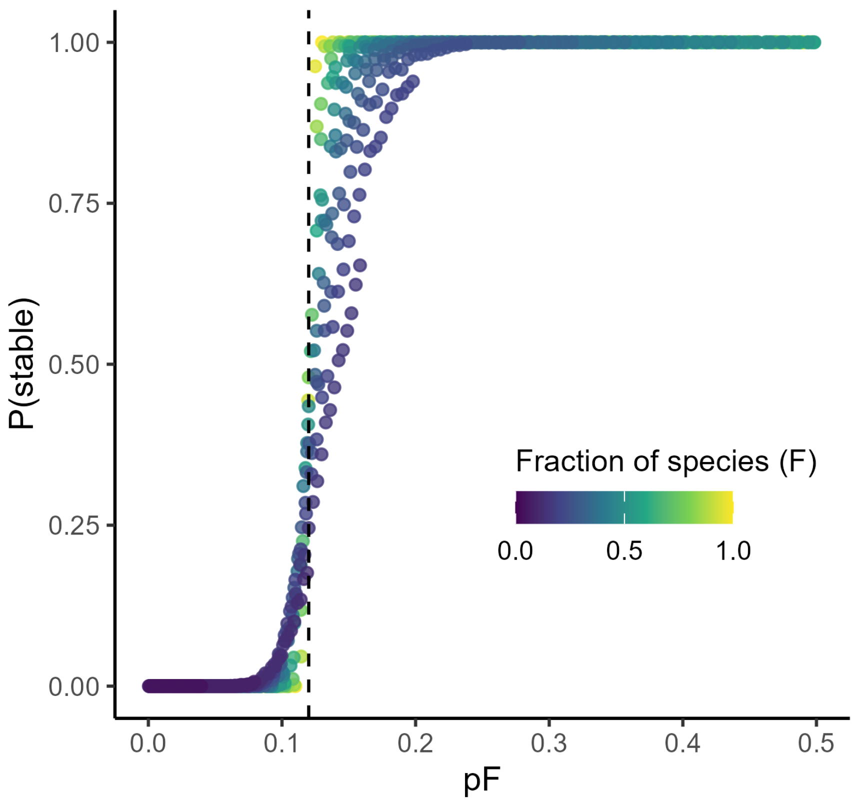

Figure 2: Threshold behavior in the probability of stabilization by dormancy when only some species go dormant. For simulations of the donor-controlled regime shown in Fig. 4 (Main Text), we plot the probability of stability (across 25 random species-subsets and 100 random food web realizations) as function of $pF$, where $F$ is the fraction of species in the community that have the capacity to go dormant and $p$ is the inactivation rate for those that do. Once the product $pF$ exceeds a threshold, equal to the value of $p$ at which stability is gained in the homogeneous case $F = 1$ ($p = \mathrm{Re}_{\max}$, dashed line), the probability of stability jumps sharply from near zero to near one. The transition is somewhat sharper for larger $F$, but the product $pF$ is highly predictive of stabilization across the entire parameter space.

allow continuous variation in $p_i$ and $q_i$ by sampling these rates *iid* for each species from an exponential distribution. To facilitate comparison with Fig. 4 (Main Text), we choose the mean of these distributions as $pF$ and $qF$, so that the mean of $p_i$ and $q_i$ matches Fig. 4 for each value of $p$ and $F$ (interpreted here as the "effective" number of species with dormancy). As in Fig. 4, we fix $q = 0.1$. We see that the results are almost indistinguishable from Fig. 4, except that stabilization is somewhat more likely in the R and B regimes. This result suggests that stability is highly predictable based only on the mean (in)activation rates across the web, despite substantial variation in these rates.

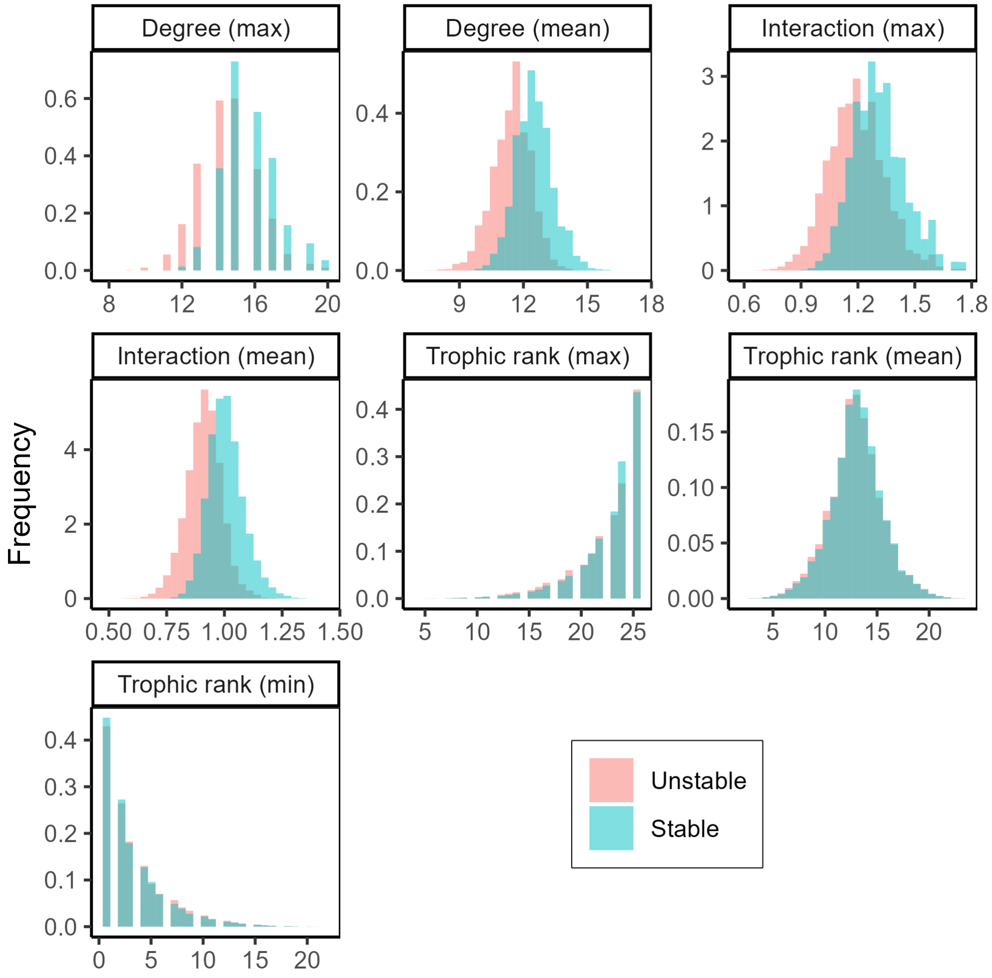

Figure 3: Conditional distribution of subset attributes by stability in the donor-controlled (D) regime. We plot the distribution of each attribute of the subset of species with dormancy over all webs in all $p - F$ combinations where the probability of stability is more than 25% but less than 75%. See text for details on subset attributes.

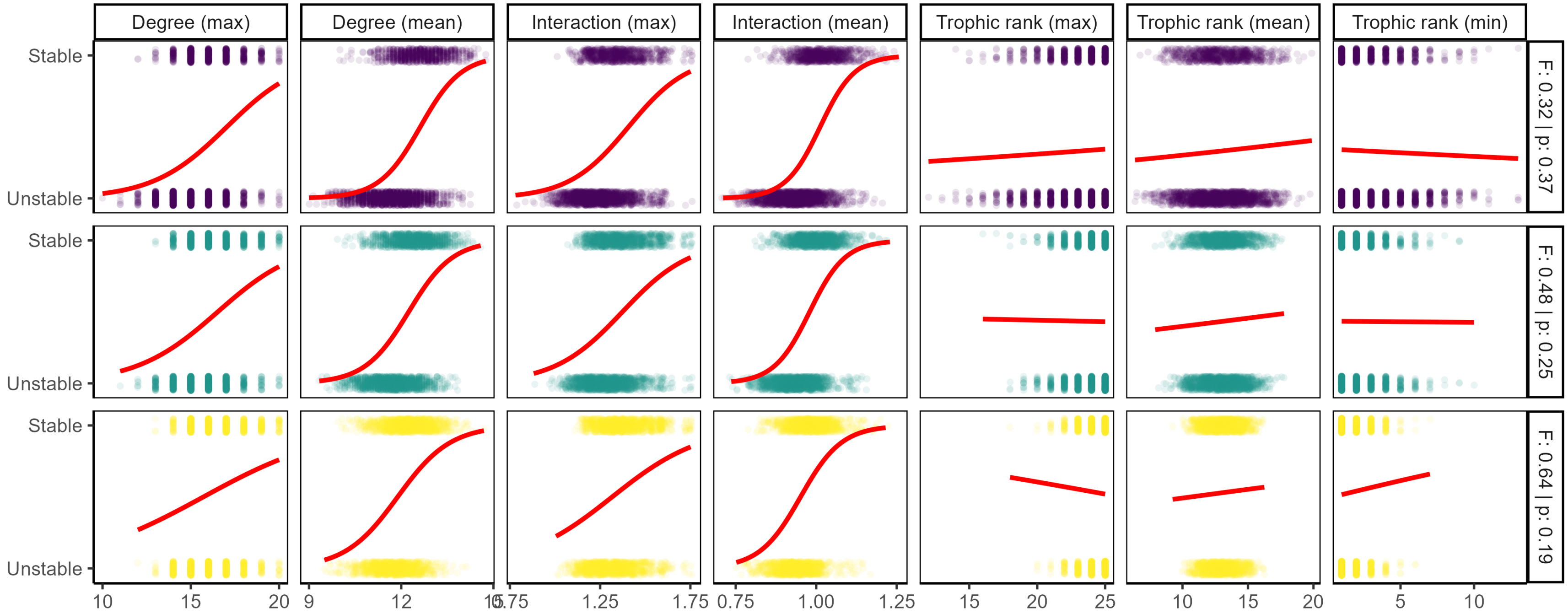

Figure 4: Stability outcomes versus subset attributes for all web-subset combinations in three $p - F$ parameter combinations along the curve $pF = \mathrm{Re}_{\max}$. See text for details on subset attributes. Red lines show logistic regressions fit using the `geom_smooth` function from the `ggplot2` package in R.

### 4 Stabilization of intransitive competition

While we focus primarily on trophic interactions – and we show that there are classes of competitive networks where dormancy can never be stabilizing (i.e., symmetrizable interaction matrices including all 2-species systems) – there are certain cases where competitive communities lose stability through Hopf bifurcations, and therefore can be stabilized by dormancy.

Consider, for example an intransitive "rock-paper-scissors" community with generalized Lotka-Volterra (GLV) dynamics [13]. We have

$$\frac{dx_i}{dt} = x_i\Big(r_i + \sum_{i=1}^{n} a_{ij}x_j\Big)\,. \tag{31}$$

with parameters $r_i = r > 0$ for all $i$ and

$$A = (a_{ij}) = \begin{pmatrix} S & \alpha & \beta \\ \beta & S & \alpha \\ \alpha & \beta & S \end{pmatrix} \tag{32}$$

with $\alpha, \beta, S \leq 0$. We show in Fig. 9 that this system can exhibit oscillatory instability associated with a Hopf bifurcation. As a result, the dynamics can be stabilized by dormancy.

This rock-paper-scissors community has an equilibrium where all species have equal abundances, so the community matrix at this equilibrium is proportional to $A$. This matrix is *circulant* (meaning that each row is identical to the preceding row, but shifted one element to the right) with well-known eigenvalues:

$$\lambda_0 = S + \alpha + \beta \qquad \lambda_{1,2} = S - \frac{1}{2}(\alpha+\beta) \pm \frac{\sqrt{3}}{2}(\alpha-\beta)i\,. \tag{33}$$

Thus, if $\alpha + \beta < 2S$, then the equilibrium is unstable, and stability is controlled by a pair of complex eigenvalues [13]. The imaginary part of these eigenvalues is proportional to $\alpha - \beta$, so dormancy can be stabilizing when each competitive interaction is sufficiently asymmetrical.

While this example shows that dormancy can play a stabilizing role in some competitive communities, we expect this effect to require very specific parameter combinations, especially in larger ecosystems. To illustrate why, we briefly consider two extensions of the intransitive competition model to include more species.

There are many ways we might generalize Eqs. 31-32 to include $n$ species. Perhaps the simplest is to consider a chain architecture where each species $i$ only interacts with $i-1$ and $i+1$ (modulo $n$). We have the interaction matrix

$$A = (a_{ij}) = \begin{pmatrix} S & \alpha & 0 & \dots & 0 & \beta \\ \beta & S & \alpha & 0 & \dots & 0 \\ 0 & \beta & S & \alpha & 0 & \dots \\ 0 & 0 & \beta & S & \alpha & \ddots \\ 0 & \ddots & \ddots & \ddots & \ddots & \ddots \\ \alpha & 0 & \dots & & \beta & S \end{pmatrix}. \tag{34}$$

This is another circulant matrix, with known eigenvalues. The spectrum of $A$ is particularly simple if we assume $\alpha < 0$ and $\beta = 0$. In this case, the eigenvalues are roots of unity (i.e., $\omega_k = \exp(2\pi ki/n)$ for $k = 0, 1, \dots, n-1$) scaled by $\alpha$ and shifted by $S$. Geometrically, these eigenvalues are equally spaced around a circle in the complex plane. For $n$ even, the rightmost eigenvalue (equal to $S - \alpha$) is real, and dormancy can never be stabilizing. For $n$ odd, the rightmost eigenvalues are complex. However, as $n$ increases, these eigenvalues become closer and closer to the real line. This implies that stabilization by dormancy is unlikely for larger $n$ (Fig. 10). For $\beta < 0$, the eigenvalues fall on a vertically-compressed ellipse, making stabilization by dormancy even more difficult.

Another simple model for $n$-species intransitive competition is one where each species "beats" and is beaten by exactly $(n-1)/2$ others (note that this requires $n$ odd). Assuming without loss of generality that $\alpha < \beta$, species $i$ "beats" species $j$ if $a_{ij} = \beta$ and $a_{ji} = \alpha$. This type of model generalizes rock-paper-scissors

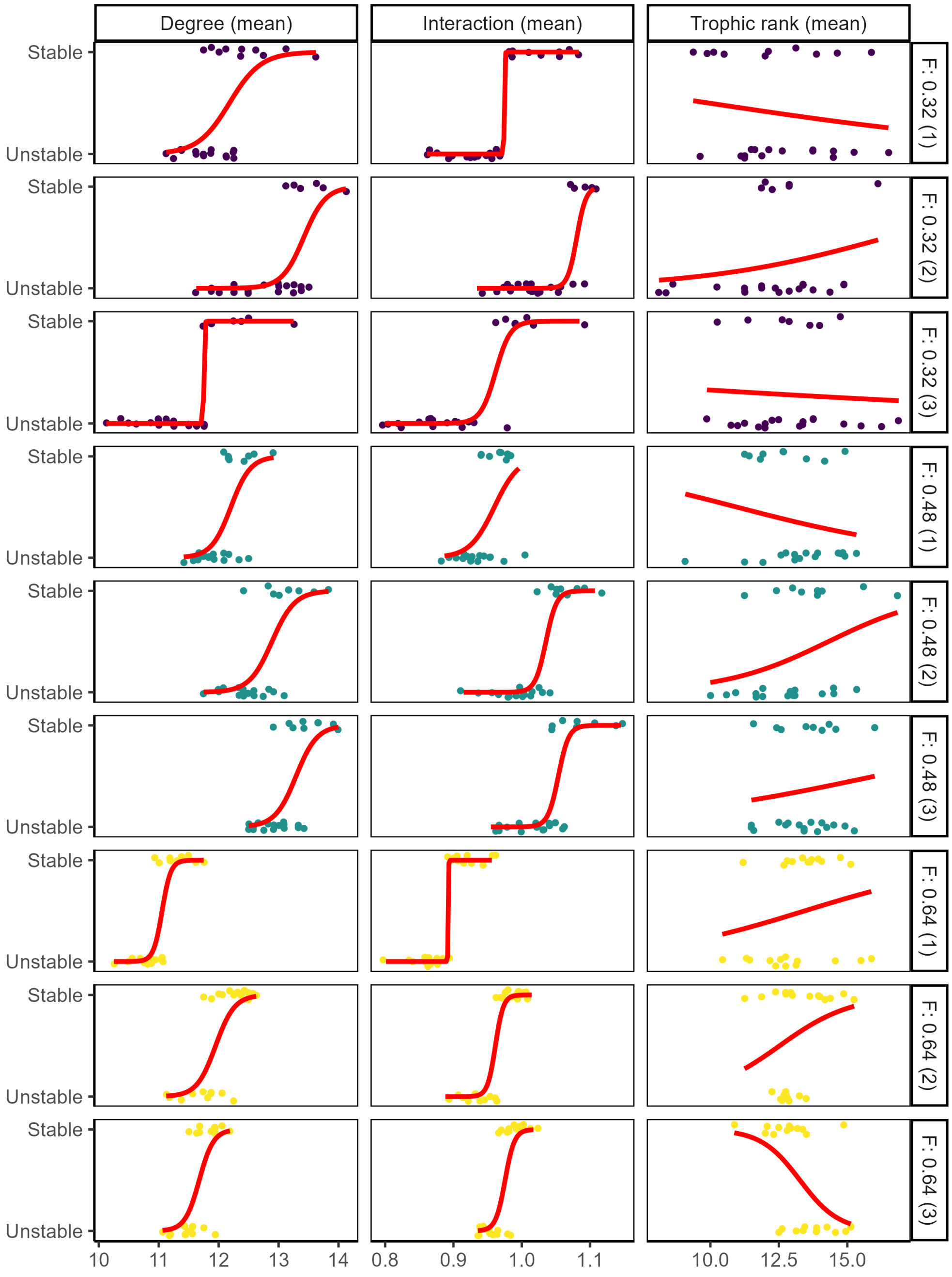

Figure 5: Stability outcomes versus subset attributes for three randomly-chosen webs (i.e., community matrix realizations) in three $p - F$ parameter combinations along the curve $pF = \text{Re}_{\text{max}}$. See text for details on subset attributes. Red lines show logistic regressions fit using the `geom_smooth` function from the `ggplot2` package in R.

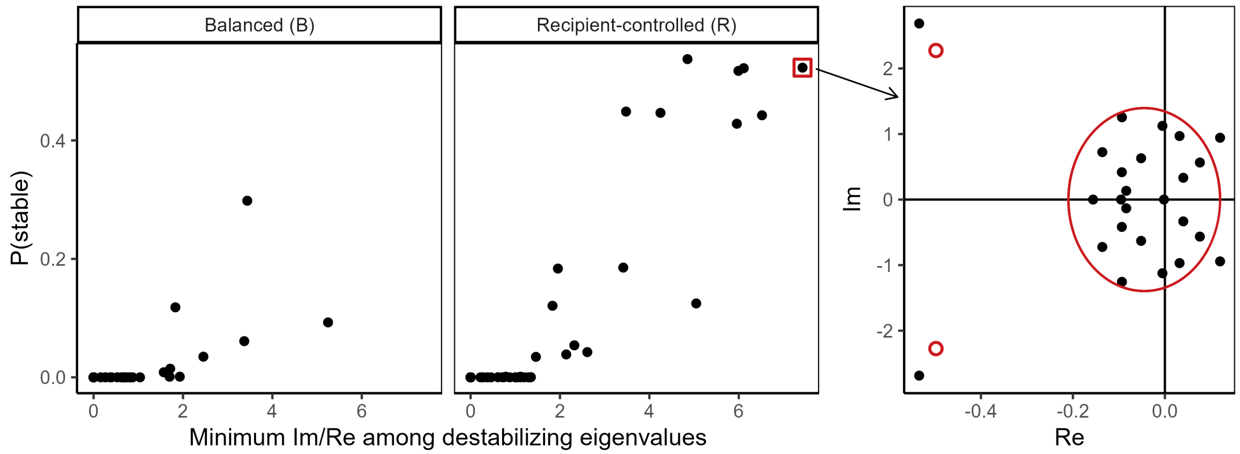

Figure 6: Probability of stability (across all $F-p$ combinations and dormant-species subsets) versus Im/Re ratio in the B and R regimes. Each point represents one food web (i.e., community matrix realization). Along the x-axis, we plot the minimum ratio of imaginary (Im) to real (Re) parts of the destabilizing eigenvalues for each web. This quantity captures how close any of the destabilizing eigenvalues (those with positive real part) fall to the real line. In particular, a value of zero indicates the presence of at least one purely real, positive eigenvalue. We find non-negligible probability of stabilization only in food webs that have large values of this quantity – i.e., in food webs whose destabilizing eigenvalues all have relatively large imaginary parts. Righthand panel shows the eigenvalues for one of these outlier food webs. By chance, none of the destabilizing eigenvalues in the bulk fall near the real line, allowing this network to be stabilized by dormancy with relatively high probability. Theoretical eigenvalue distribution in red.

to "rock-paper-scissors-lizard-Spock" and so on. We have the interaction matrix

$$A = (a_{ij}) = \begin{pmatrix} S & \alpha & \alpha & \dots & \beta & \beta \\ \beta & S & \alpha & \alpha & \dots & \beta \\ \beta & \beta & S & \alpha & \alpha & \dots \\ \vdots & \beta & \beta & S & \alpha & \ddots \\ \beta & \ddots & \ddots & \ddots & \ddots & \ddots \\ \alpha & \alpha & \dots & \beta & \beta & S \end{pmatrix} \tag{35}$$

which is once again circulant. We can easily characterize the eigenvalues of this interaction matrix. In general, a circulant matrix whose first row is $(a_0, a_1, \dots, a_{n-1})$ has eigenvalues

$$\lambda_k = \sum_{j=0}^{n-1} a_j \, \omega_k^j \,, \quad k = 0, 1, \dots, n-1 \tag{36}$$

where $\omega_k = \exp(2\pi k i/n) = \cos(2\pi k/n) + i\sin(2\pi k/n)$ are again the $n$th roots of unity. Applying this formula to Eq. 35, we find that

$$\lambda_k = S + \sum_{j=1}^{(n-1)/2} \alpha \, \omega_k^j + \beta \, \omega_k^{n-j} = S + \sum_{j=1}^{(n-1)/2} \alpha \, \omega_k^j + \beta \, \omega_k^{-j} \tag{37}$$

where the last equivalence follows from the fact that $\omega_k^n = 1$. One eigenvalue, corresponding to $k = 0$ is easy to see: we have $\omega_0 = 1$, so $\lambda_0 = S + \sum_{j=1}^{(n-1)/2} \alpha + \beta = S + (n-1)(a+b)/2$. This eigenvalue is real and negative. To characterize the remaining eigenvalues, we write real and imaginary parts separately. Using

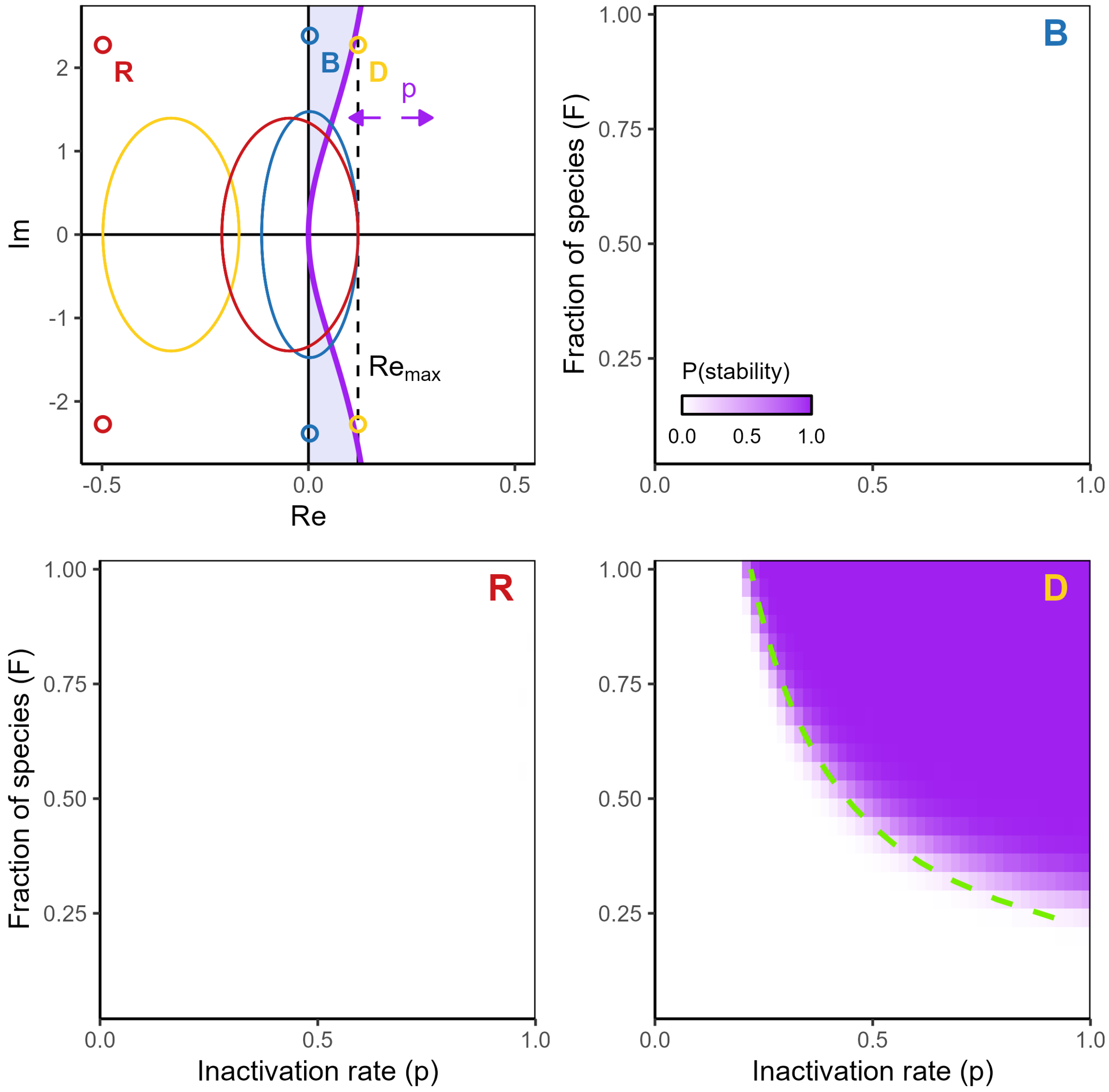

Figure 7: Dormancy stabilizes structured food webs when only some species go dormant. As Fig. 4 (Main Text) except that $q = 2$. In the top left panel, we plot the ASD for $p = 0.2$. Note that the (theoretical) leading outlier eigenvalue in the $D$ regime is still not stabilized at this value of $p > \mathrm{Re}_{\max}$, a consequence of larger $q$ pushing the asymptotic phase of the ASD farther from the real axis. However, as in Fig. 4, once the product $pF$ exceeds the value of $p$ where stabilization occurs in the homogeneous case (here, $pF \approx 0.22$; green dashed line), stabilization is very likely in the $D$ regime. In the other two regimes, the increase in $q$ makes stabilization much less likely overall. Panels $B$ and $R$ appear blank because stabilization is very rare in these regimes; in the $B$ regime, we never find stabilization by dormancy across all simulations, and in the $R$ regime, the probability of stabilization is less than 3% for all $p - F$ combinations.

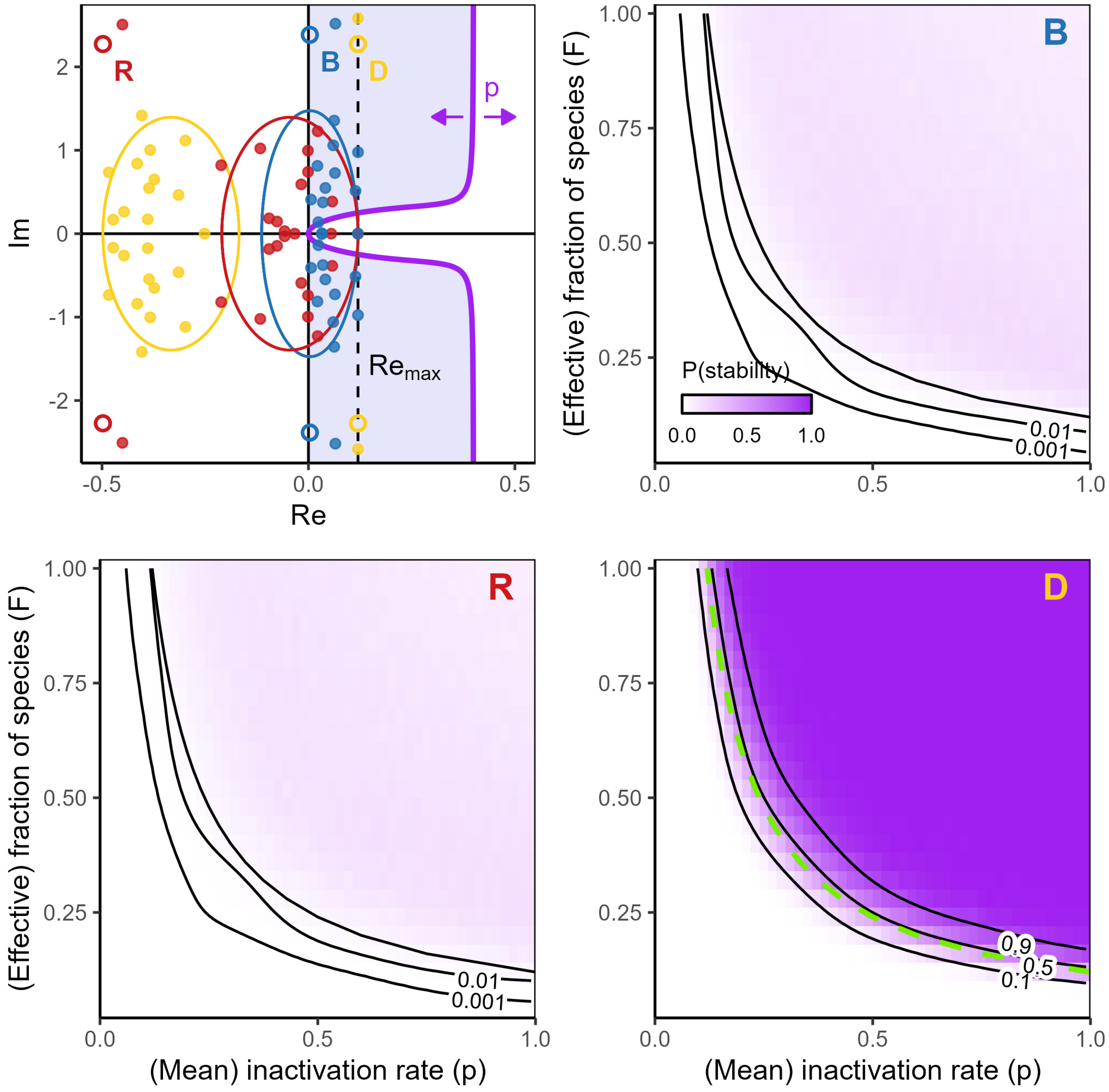

Figure 8: Dormancy stabilizes structured food webs when species enter and exit dormancy at different rates. As Fig. 4 (Main Text) except that all $p_i$ and $q_i$ are sampled *iid* from exponential distributions, as described in the text.

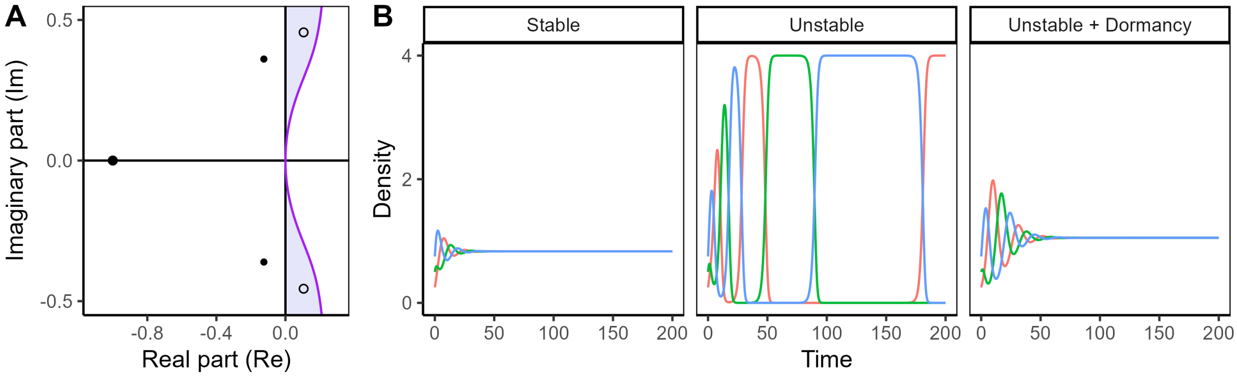

Figure 9: Dormancy stabilizes “rock-paper-scissors” competition. Eigenvalues (A) and dynamics (B) for Eq. 32 with $r = 1$, $\alpha = -0.6$, and $\beta = -0.1$. With $S = -0.5$, the dynamics are stable (filled points in panel A), with $S = -0.25$ (open points in panel A) the system loses stability through a Hopf bifurcation. The dynamics approach a heteroclinic cycle, jumping abruptly between periods of dominance for each species (colors), with the other two reaching very low densities [13]. With dormancy, these dynamics are stabilized, and the system settles back into equilibrium. Dormancy parameters: $p = 0.25$, $q = 0.2$.

Euler's formula, $\omega_k^j = \exp(2\pi jki/n) = \cos(2\pi jk/n) + i\sin(2\pi jk/n)$, so we have

$$\mathrm{Re}(\lambda_k) = S + (a+b) \sum_{j=1}^{(n-1)/2} \cos(2\pi jk/n) \tag{38}$$

using the fact that the cosine is an even function. To compute the sum, we apply Lagrange's identity, yielding

$$\mathrm{Re}(\lambda_k) = S + (a+b)\frac{-\sin(\pi k/n) + \sin(\pi k)}{2\sin(\pi k/n)} = S + (a+b)\frac{-\sin(\pi k/n)}{2\sin(\pi k/n)} = S - \frac{a+b}{2}\,. \tag{39}$$

The dependence on $k$ disappears, indicating that the remaining $n-1$ eigenvalues lie on a vertical line in the complex plane.

Now we consider their imaginary parts:

$$\mathrm{Im}(\lambda_k) = (b-a) \sum_{j=1}^{(n-1)/2} \sin(2\pi jk/n) \tag{40}$$

using the fact that the sine is an odd function. Again, we apply Lagrange's identity to simplify the sum:

$$\mathrm{Im}(\lambda_k) = (b-a)\frac{\cos(\pi k/n) - \cos(\pi k)}{2\sin(\pi k/n)} = (b-a)\frac{\cos(\pi k/n) \pm 1}{2\sin(\pi k/n)}. \tag{41}$$

Consider the eigenvalue(s) corresponding to $k = 2$ (and $(n-3)/2$). For sufficiently large $n$, we can apply small-angle approximations to write $\cos(2\pi/n) \approx 1 - \frac{1}{2}(2\pi/n)^2$ and $\sin(2\pi/n) \approx 2\pi/n$. So $\mathrm{Im}(\lambda_2) \approx \frac{1}{2}(\pi/n)$. We see that the imaginary part of this eigenvalue is proportional to $1/n$. Thus, we find once again that some leading eigenvalues have small real parts for $n$ large (Fig. 11).

These examples are highly symmetrical, and they introduce long intransitive loops, which may drive low frequency oscillations associated with eigenvalues with small imaginary parts. Other intransitive architectures are possible, for example with small intransitive loops nested inside larger ones. Further investigation of more complex intransitive networks is beyond the scope of our study, but the examples above suggest that it may be challenging to find large networks that are stabilized by dormancy.

#### 5 Additional parameter and simulation details

In Fig. 3 (Main Text), we generate time-series corresponding to scenarios with and without dormancy, in order to visualize the effect of dormancy on stability. This requires us to specify a dynamical model (while

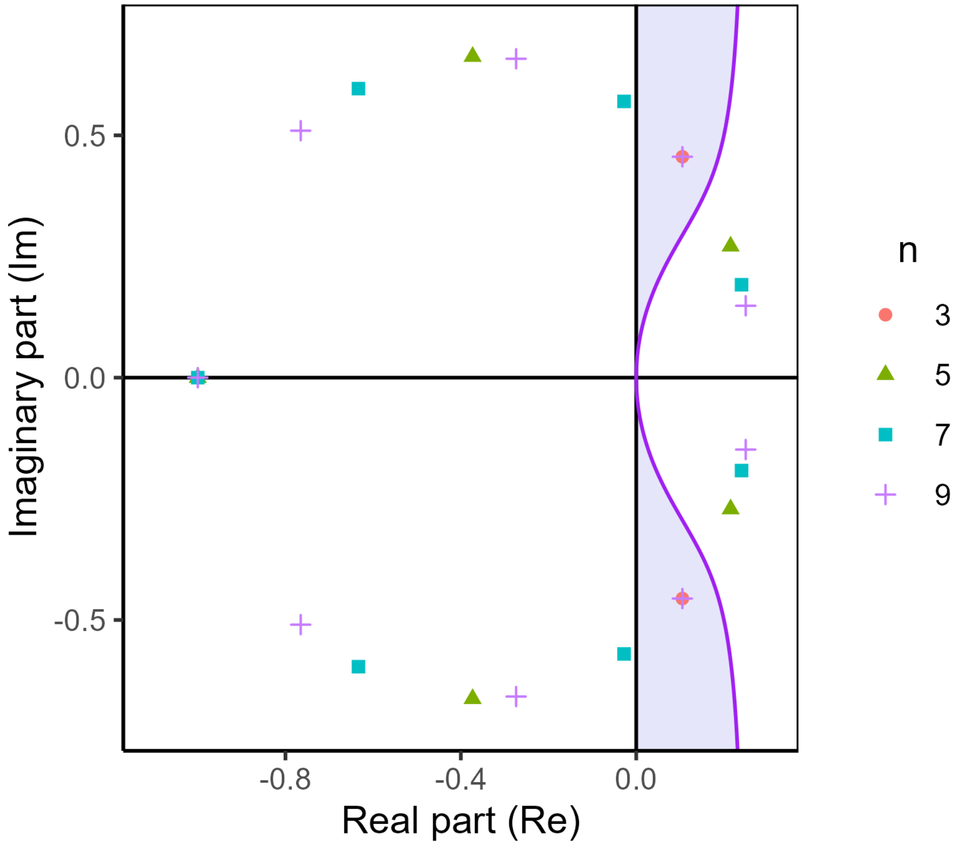

Figure 10: Eigenvalues for an intransitive chain (Eq. 34) with $n$ species. Here, $r = 1$, $\alpha = -0.6$, $\beta = -0.1$, and $S = -0.25$, as in Fig 9. For each $n$, eigenvalues lie on a circle in the complex plane; as $n$ grows, the leading eigenvalues are closer to the real line. While dormancy is stabilizing for the three-species system, for $n \geq 5$, it becomes insufficient to stabilize the dynamics. Note that only odd values of $n$ are shown because chains with even $n$ have a real destabilizing eigenvalue (see text). Dormancy parameters: $p = 0.25$, $q = 0.2$.

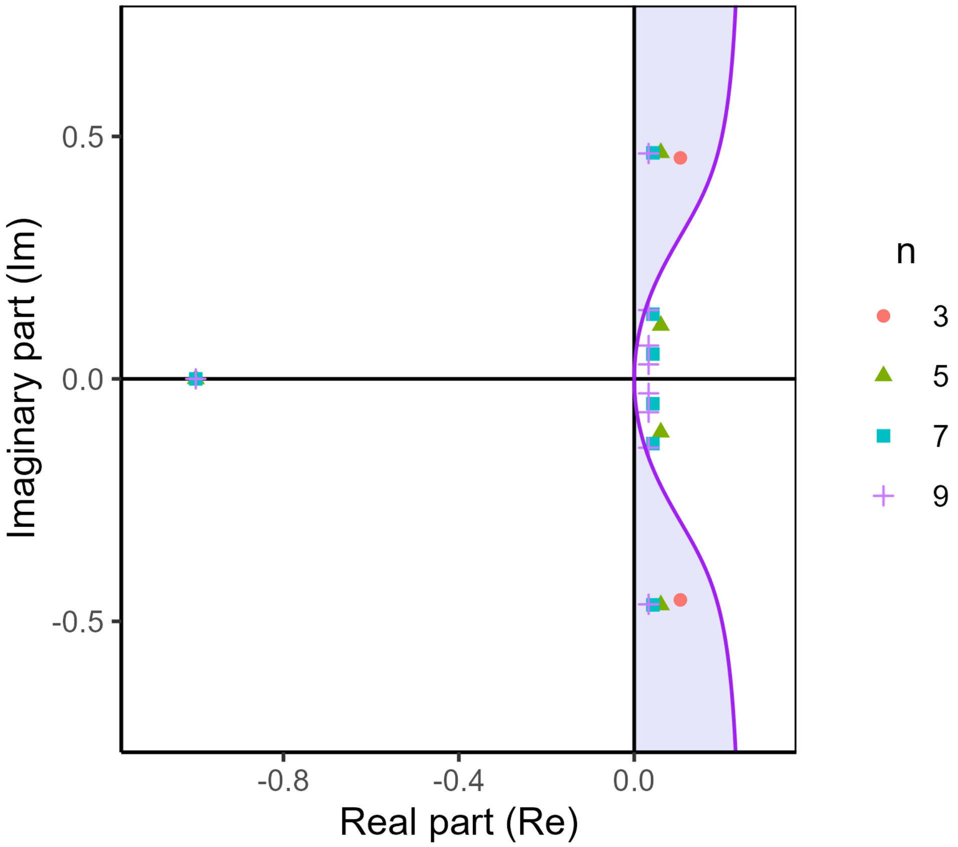

Figure 11: Eigenvalues for generalized rock-paper-scissors competition (Eq. 35) with $n$ species. Here, $r = 1$, $\alpha = -0.6$, $\beta = -0.1$, and $S = -0.25$, as in Fig 9. For each $n$, all eigenvalues except one lie on a vertical line in the complex plane; as $n$ grows, some of these (leading) eigenvalues become closer to the real line. While dormancy is stabilizing for the three-species system, for $n \geq 5$, it becomes insufficient to stabilize the dynamics. Note that the location of the line of eigenvalues changes slightly between different values of $n$ because the community matrix is scaled by equilibrium abundances (which decrease with $n$). Dormancy parameters: $p = 0.25$, $q = 0.2$.

the community matrices could correspond to many alternative models). For simplicity, we use generalized Lotka-Volterra (GLV) models to generate the dynamics. The GLV model dynamics are defined by:

$$\frac{dx_i}{dt} = x_i\Big(r_i + \sum_{i=1}^{n} a_{ij}x_j\Big)\,. \tag{42}$$

This model is somewhat unrealistic for trophic interactions, corresponding to assuming Type I (linear) functional responses. However for the purposes of illustration, this choice is convenient, because it allows us to easily select parameters of the dynamical model that lead to the desired community matrix. The local dynamics around the equilibrium point will be the same for more complex models with more realistic functional responses.

To parameterize the GLV model given a target community matrix, $M$ (generated by one of our four community matrix models), we set $A = M$ and "plant" an equilibrium with $x_i^\star = 1$ for all $i$ by setting $\mathbf{r} = -A\mathbf{x}^\star$. The community matrix for the GLV model at $\mathbf{x}^\star$ is $D(\mathbf{x}^\star)\mathbf{A} = IM = M$, as desired.

We used the following parameter choices to generate example community matrices in Fig. 3:

- **Panel A:** $n = 50$; elements $m_{ij}$ sampled independently from a normal distribution with $\mu = 0$, $\sigma^2 = 1/n = 0.02$; self-regulation $S = 0.6$.
- **Panel B:** $n = 50$; magnitudes $|m_{ij}|$ sampled independently from a half-normal distribution with $\sigma^2 = 1/n = 0.02$; self-regulation $S = 0.2$.
- **Panel C:** $n = 9$, $l = 4.5$, $u = 5$, $S = 0.5$, $s_a = -3.5$, $s_b = -0.1$. Note that these parameter choices lead to $x \approx -0.02$ and $y \approx -0.74$ in Eq. 3, consistent with our approximation assumption that $|x|, |y| < 1$.
- **Panel D:** $n = 50$; upper-triangular elements sampled independently from a half-normal distribution with $\sigma^2 = 0.04/n$, leading to $\mu_U = \sigma\sqrt{2/\pi} \approx 0.0225$, $\sigma_U^2 = (0.04/n)(1-2/\pi) \approx 2.9 \times 10^{-4}$, lower-triangular elements sampled independently from a half-normal distribution with $\sigma^2 = 1/n$, leading to $\mu_L = \sigma\sqrt{2/\pi} \approx 0.113$, $\sigma_L^2 = (1/n)(1-\pi/2) \approx 0.007$; self-regulation $S = 0.4$.

In Fig. 4, we sample elements of $M$ from a Gamma distribution in order to control the mean and variance separately. We also use incomplete connectance ($C = 0.5$). The three regimes considered in the Main Text are defined by sampling nonzero interactions from Gamma distributions with the following statistics:

- **Donor-controlled (D) regime:** $\mu_U = 0.4$, $\sigma_U^2 = 0.01$, $\mu_L = 0.2$, $\sigma_L^2 = 0.0025$.
- **Balanced-interaction (B) regime:** $\mu_U = 0.3$, $\sigma_U^2 = 0.005625$, $\mu_L = 0.3$, $\sigma_L^2 = 0.005625$.
- **Recipient-controlled (R) regime:** $\mu_U = 0.2$, $\sigma_U^2 = 0.0025$, $\mu_L = 0.4$, $\sigma_L^2 = 0.01$.
